# Ubiquitin-dependent signal amplification in lipid saturation sensing

**DOI:** 10.1101/2025.11.05.686737

**Authors:** Jona Causemann, Barbara Schmidt, Daniel Granz, Thorsten Mosler, Martin Jung, Ivan Dikic, Heiko Rieger, Robert Ernst

## Abstract

Cellular membranes are dynamic platforms whose composition and biophysical properties are surveyed by sensor proteins to maintain homeostasis. Failure to preserve membrane homeostasis, however, results in cellular stress and organelle dysfunction. Using the prototypical lipid saturation sensor Mga2, we explored how weak physical cues that modulate rotational movements in the transmembrane region are converted into decisive biochemical outputs that ultimately control the production of unsaturated fatty acids. Quantitative *in vitro* ubiquitylation assays and kinetic modeling reveal vastly distinct rates of Mga2 ubiquitylation controlled by the membrane environment. Mga2 ubiquitylation dominates in tightly packed, saturated membranes, while loosely packed environments favor an inhibitory autoubiquitylation of the cognate E3 ligase Rsp5. This mechanism provides a means of signal amplification, which can function even in the absence of deubiquitylating enzymes. Our findings provide a mechanistic framework for how membrane property sensors convert weak, fluctuating physical cues into robust biochemical outcomes, and put a spotlight on the regulatory potential of E3 ligase autoubiquitylation in cellular surveillance.

## Introduction

Biological membranes are dynamic structures whose physical properties — such as thickness, compressibility, surface charge, and lipid packing — are critical for cellular function (Bigay & Antonny, 2012; Cybulski *et al*, 2010; Holthuis & Menon, 2014; Renne & Ernst, 2023). Eukaryotic cells regulate these properties across subcellular compartments, ensuring that each membrane maintains its characteristic compositions and properties suited to its role (Harayama & Riezman, 2018; Renne & Ernst, 2023). This balance, known as biophysical membrane homeostasis, is maintained by sensor proteins that detect deviations in membrane properties and trigger adaptive responses with a wide impact on lipid metabolism, protein quality control, and membrane trafficking (Ernst *et al*, 2018; Covino *et al*, 2018; Harayama & Riezman, 2018).

Membrane property sensors are diverse in form and function, yet they share two defining features: they are highly sensitive to specific physical parameters of the lipid bilayer, and they are structurally metastable, allowing them to respond to subtle fluctuations in the membrane environment (Antonny, 2011; Covino *et al*, 2018). Examples include sensors of curvature elastic stress (Boumann *et al*, 2006; Haider *et al*, 2018), lipid saturation (Covino *et al*, 2016; Ballweg *et al*, 2020; Vrentzou *et al*, 2025), and reduced membrane compressibility (Halbleib *et al*, 2017; Alsayyah *et al*, 2025). Intriguingly, lipid saturation sensors have been implicated in controlling the cellular response to low temperature and hypoxia (Zhang *et al*, 1999; Jiang *et al*, 2001; Nakagawa *et al*, 2002; Jiang *et al*, 2002), while a membrane compressibility sensor coordinates the synthesis of membrane lipids and proteins via the unfolded protein response (UPR) of the endoplasmic reticulum (ER) (Renne & Ernst, 2023; Ernst *et al*, 2024). Even though the biological functions of these pathways are well established, the mechanisms by which small changes in membrane properties are translated into decisive signaling remain poorly understood.

One of the best-characterized pathways for biophysical membrane homeostasis is the OLE pathway in *Saccharomyces cerevisiae* (Hoppe *et al*, 2000; Rape *et al*, 2001; Shcherbik *et al*, 2003; Ballweg & Ernst, 2017). It controls lipid packing in the ER membrane by regulating the production of unsaturated fatty acids (UFAs) and operates through two ER-membrane–anchored transcription factors, Mga2 and Spt23, which sense membrane lipid saturation to regulate expression of the essential fatty acid desaturase gene *OLE1* (Stukey *et al*, 1989; Zhang *et al*, 1999; Hoppe *et al*, 2000; Covino *et al*, 2016). Among the two, Mga2 plays the dominant role: deletion of *MGA2* leads to reduced *OLE1* expression, accumulation of saturated lipids, and lipid bilayer stress with pronounced morphological changes of the ER membrane (Chellappa *et al*, 2001; Jiang *et al*, 2001; Surma *et al*, 2013). To become active, Mga2 and Spt23 must be released from their membrane anchor by the proteasome before they can translocate to the nucleus (Hoppe *et al*, 2000). Sensor ubiquitylation is carried out by the E3 ubiquitin ligase Rsp5 and inhibited by unsaturated lipids (Hoppe *et al*, 2000; Shcherbik *et al*, 2003). Cleavage by the proteasome at an internal site (Piwko & Jentsch, 2006) and -according to our working model-degradation of the membrane-anchor liberates the active transcription factor, which then migrates to the nucleus and upregulates *OLE1* expression (Hoppe *et al*, 2000; Rape *et al*, 2001; Ballweg & Ernst, 2017). Newly synthesized, coenzyme A-activated UFAs can be incorporated into membrane lipids to restore normal membrane lipid packing (Ernst *et al*, 2016), thereby closing the regulatory circuit.

At the center of this sense-and-respond circuit is a biophysical mechanism provided by the transmembrane helix of Mga2 (Covino *et al*, 2016). Each protomer of the dimeric protein contains a bulky tryptophan that probes lipid packing in the hydrophobic core of the ER membrane at the level of Δ9 double bonds in lipid fatty acyl chains. The dimeric transmembrane helices dynamically sample different rotational states and the overall structural organization renders Mga2 sensitive to lipid saturation, but less sensitive to the lipid headgroup composition (Covino *et al*, 2016; Ballweg *et al*, 2020). When lipid packing is high (i.e. membranes are more saturated), the tryptophan residues are more likely to face each other in the dimer interface, thereby stabilizing a spectrum of conformations that promotes Mga2 ubiquitylation. When lipid packing is low, the helices populate alternative conformations, and ubiquitylation efficacy drops (Covino *et al*, 2016; Ballweg *et al*, 2020). Notably, even modest shifts in the population of these conformations — as seen in coarse-grained molecular dynamics simulations and continuous-wave electron paramagnetic resonance (cwEPR) experiments — are sufficient to drive changes in Mga2 activity both *in vitro* and *in vivo* (Covino *et al*, 2016; Ballweg *et al*, 2020). This raises a central question: How do these rather subtle, membrane-driven conformational changes lead to decisive ubiquitylation events occurring ≈50 amino acids away, in the cytosol?

The ubiquitylation machinery that contributes to Mga2 activation follows the canonical E1–E2–E3 cascade: ubiquitin is activated by an E1 enzyme, transferred to a conjugating enzyme (E2), and finally attached to substrate lysines by an E3 ubiquitin ligase (Shcherbik *et al*, 2003; Pohl & Dikic, 2019). In the case of Mga2, the E3 is Rsp5, a HECT-domain containing ligase from the Nedd4 family with a multitude of cellular functions (Shcherbik *et al*, 2004; Bhattacharya *et al*, 2009, 2008). Rsp5 contains an N-terminal C2 domain, three WW domains for substrate recognition, and a catalytic HECT domain. While the third WW domain and the HECT domain are sufficient to complement *RSP5*’s essential functions (Hoppe *et al*, 2000), the roles of the N-terminal C2 and the other two WW domains can vary depending on the pathway and substrate (Wang *et al*, 1999; Chang *et al*, 2000; Dunn & Hicke, 2001; Bhattacharya *et al*, 2008). Like other HECT domain ligases, Rsp5 is thought to transfer ubiquitin sequentially, building polyubiquitin chains on its client one molecule at a time (Wang & Pickart, 2005; Pierce *et al*, 2009; French *et al*, 2017). Apart from its role in lipid regulation, Rsp5 is involved in endocytosis, the heat shock response, and other stress-related pathways — but how it allocates its activity among these competing demands remains unclear (Kaliszewski & Zoładek, 2008; Sardana & Emr, 2021). Interestingly, Rsp5 can autoubiquitylate and dampen its own ligase activity (Attali *et al*, 2017).

Ubiquitylation is a remarkably versatile regulatory tool. The anaphase-promoting complex (APC), for instance, establishes a defined temporal order of degradation for a variety of key cell-cycle regulators in G1 and mitosis by sequential ubiquitylation of its substrates (Peters, 2002; Rape *et al*, 2006). Pioneering work has demonstrated that the processivity of multiubiquitylation in conjunction with deubiquitylating enzymes (DUBs) determines this ordering of substrates (Peters, 2002). Similarly, the ER-associated degradation (ERAD) depends on the selective ubiquitylation of membrane and lumenal client proteins, which targets them for proteasomal clearance (Needham *et al*, 2019; Christianson & Carvalho, 2022). Reconstituting the decision-making process revealed that a deubiquitylating activity is critical for amplifying modest differences in E3 ligase-substrate interactions to facilitate robust decisions (Zhang *et al*, 2013). These systems showcased how E3 ligase processivity combined with deubiquitylating activities can translate weak or transient signals into all-or-none decisions by amplifying small molecular changes (Rape *et al*, 2006; Zhang *et al*, 2013).

Inspired by this principle, we wondered whether biophysical changes in the lipid environment of Mga2 can be amplified by the coupled ubiquitylation reaction even in the absence of a DUB, when negative feedback is supplied by Rsp5’s autoubiquitylation, which reduces its activity and limits signal propagation (Attali *et al*, 2017). This would establish an additional mode of decision-making that deviates from classical signaling pathways with push-pull architectures (e.g. ubiquitylation/DUB or kinase/phosphatase cycles) without, however, excluding a contribution of DUBs to the OLE pathway (Ferrell & Xiong, 2001; Markevich *et al*, 2004; Rumpf & Jentsch, 2006; Stegmeier *et al*, 2007; Zhang *et al*, 2013; Ballweg & Ernst, 2017; Hansen *et al*, 2019). We set out to test whether the dynamic interplay of the E3 ligase with its client alone is sufficient to amplify small differences in membrane lipid packing. Using a reconstituted *in vitro* system with purified components and fluorescent in-gel readouts, we quantified Mga2 ubiquitylation and modeled its kinetics. Even though the recruitment of Rsp5 to Mga2 is independent of the lipid environment initially, we find that signal amplification is provided by the first five ubiquitin transfers, yielding up to 4.5-fold higher levels of ubiquitylated Mga2 species in a more saturated membrane environment required for transcription factor mobilization. Strikingly, the lipid environment of Mga2 also provides negative feedback by diverting the ubiquitylation toward inhibitory Rsp5 autoubiquitylation in loosely packed membranes. Together, our findings support a signal amplification mechanism in which a combination of positive feedback and E3 ligase autoinhibition serve as regulatory logic. This mechanism may represent a more general strategy by which cells translate subtle biophysical inputs into decisive molecular responses.

## Results

### Development of a fully defined *in vitro* assay to study Mga2 ubiquitylation

Using the lipid saturation sensor Mga2 as a paradigm, we wanted to learn how fluctuating signals from the transmembrane region (Covino *et al*, 2016; Ballweg *et al*, 2020) can be amplified to robustly regulate transcription factor activation. Suspecting an important contribution of sensor ubiquitylation, we first optimized the reconstitution of the minimal sense-and-response construct ^ZIP-MBP^Mga2 in liposomes (Ballweg *et al*, 2020). The construct contains an N-terminal leucine zipper (ZIP) for efficient dimerization, the maltose binding protein (MBP) from *Escherichia coli* as an affinity and solubility tag, and both the juxtamembrane (aa 951-1037) and C-terminal transmembrane region (aa 1038-1055) of Mga2 (Figure 1A). The juxtamembrane region includes the ^967^LPKY^970^ motif for recruiting the E3 ubiquitin ligase Rsp5, and three, partially redundant lysine residues (K980, K983, K985) that are ubiquitylated by Rsp5 *in vivo* (Shcherbik *et al*, 2004; Bhattacharya *et al*, 2009). We installed a cysteine at position S1003C for fluorescence labeling using ATTO488-maleimide to facilitate in-gel quantification of ubiquitylated ^ZIP-MBP^Mga2 with a broad dynamic range. After purifying labeled ^ZIP-MBP^Mga2 in octyl-β-D-glucopyranoside (OG) (Ballweg *et al*, 2020), we reconstituted the sensor protein in liposomes at a molar protein-to-lipid ratio of 1:8000. Quantitative detergent removal was accomplished by dialysis against a detergent-free buffer containing SM-2 BioBeads (BBs). We used a 1-palmitoyl-2-oleoyl-*sn*-glycero-3-phosphocholine (POPC) lipid matrix to mimic conditions that facilitate Mga2 ubiquitylation, and a loosely packed 1,2-dioleoyl-*sn*-glycero-3-phosphocholine (DOPC) matrix to mimic conditions that do not support transcription factor modification and activation (Covino *et al*, 2016; Ballweg *et al*, 2020). Labeled ^ZIP-MBP^Mga2 was successfully reconstituted in both lipid environments with protein recoveries of >70% (Figure 1B) yielding proteoliposomes with an average diameter of ≈80 nm as studied by dynamic light scattering (DLS) (Figure 1C). In contrast to previous reconstitution protocols (Ballweg *et al*, 2020), our optimized procedures resulted in an almost unidirectional topology of the reconstituted protein as judged from proteinase K protection assays with >80% of the sequence elements relevant for ubiquitylation being accessible from the outside of the proteoliposomes (Figure 1D). The successful reconstitution was further validated by sucrose density gradient centrifugation (Supplementary Figure S1A), and extraction assays using sodium carbonate, urea, and high salt (Supplementary Figure S1B). Hence, the sensor protein was successfully and stably integrated into the lipid bilayer.

**Figure 1:**
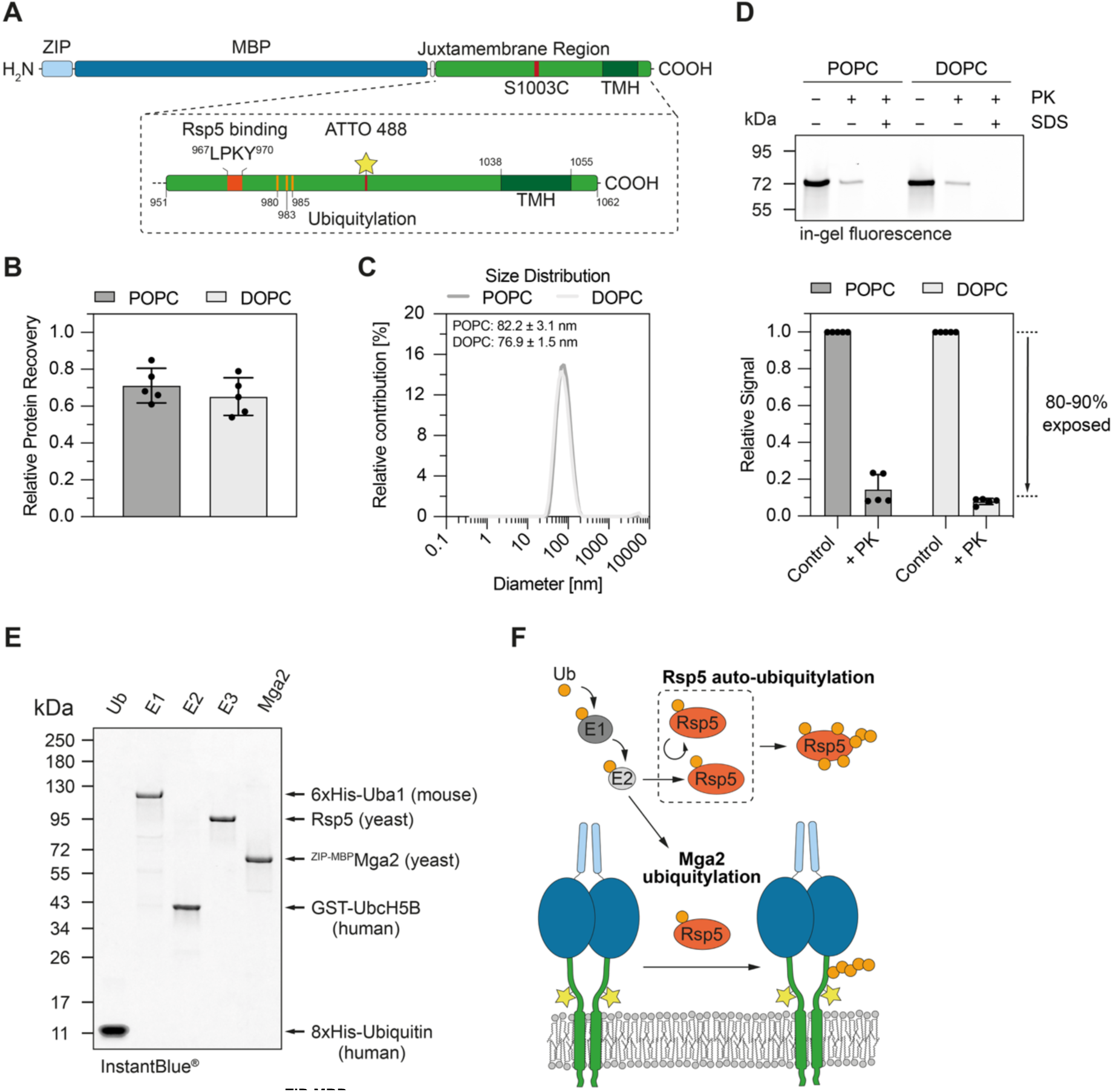
Reconstitution of ^ZIP-MBP^Mga2 in distinct lipid environments. **(A)** Schematic representation of the Mga2 construct comprising an N-terminal leucine zipper (ZIP), a maltose binding protein (MBP), and a Mga2-derived sequence (aa 951-1062). The juxtamembrane region of Mga2 (aa 951-1037) contains a binding site for the E3 ubiquitin ligase Rsp5 (^967^LPKY^970^) and three lysine residues used in vivo for ubiquitylation (K980, K983, K985). Serine 1003 found in wild-type Mga2 is replaced by cysteine (S1003C) for ATTO 488 labeling and fluorescence detection. Mga2’s transmembrane helix (TMH, Mga2 aa 1038-1055) anchors it in the membrane. (**B)** ^ZIP-MBP^Mga2 recovery monitored by ATTO 488 fluorescence after reconstitution in liposomes. Proteoliposomes were solubilized with 75 mM OG to reduce scattering. Background-corrected intensities were normalized to the intensity of the input sample prior to the reconstitution. The mean and SD of n = 5 independent reconstitution experiments are plotted.**(C)** Size of proteoliposomes measured by dynamic light scattering, showing mean diameters for POPC and DOPC environments (n = 3 reconstitutions). **(D)** Proteinase K protection assay to assess ^ZIP-MBP^Mga2 orientation in proteoliposomes. Samples were separated by SDS-PAGE and analyzed by in-gel fluorescence scanning. Graphs show the mean and SD of n = 5 independent experiments. **(E)** SDS-PAGE analysis of recombinant proteins (0.5 µg per lane) used for the *in vitro* ubiquitylation of ^ZIP-MBP^Mga2, stained with InstantBlue^®^. **(F)** Scheme of Mga2 *in vitro* ubiquitylation, detailing the roles of the E1, E2, and E3 enzymes, as well as highlighting Rsp5 autoubiquitylation.

**Figure EV1:**
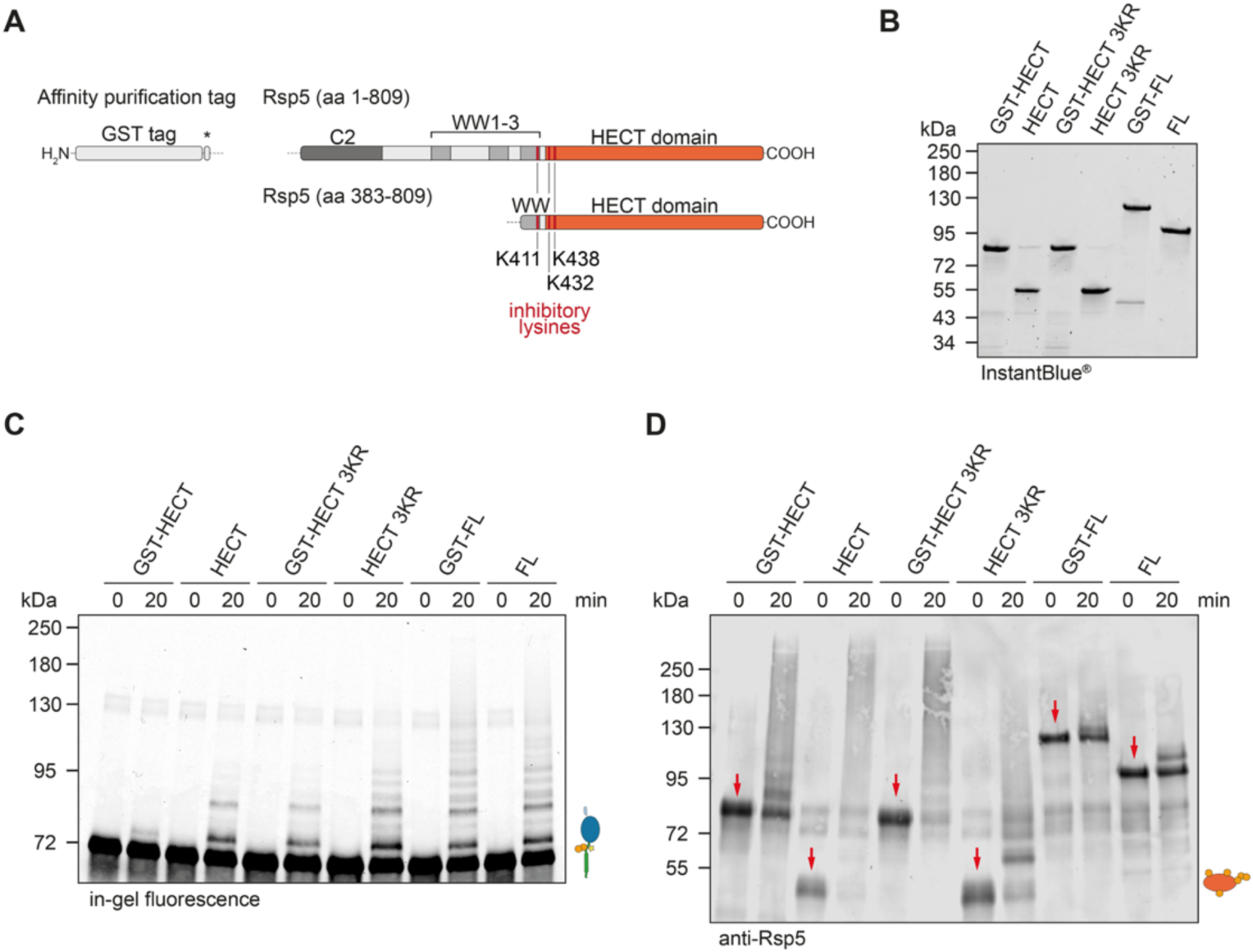
Full-length Rsp5 supports ^ZIP-MBP^Mga2 ubiquitylation *in vitro*. **(A)** Domain architecture of full-length Rsp5 (aa 1-809) and a truncated version (aa383-809). The full-length construct contains an N-terminal C2 domain (dark gray), three WW domains (gray), and the catalytic HECT domain (orange). The short construct encompasses only the third WW and the HECT domain. Both constructs are purified via an N-terminal GST tag that can be removed by protease cleavage using a TEV (truncated construct) or HRV 3C protease (full-length construct). Three lysine residues that inhibit the E3 ligase activity upon ubiquitylation are highlighted in red. **(B)** SDS-PAGE of recombinant Rsp5 construct after isolation from *E. coli*. GST-HECT: truncated construct (aa 383-809) containing the affinity purification tag. HECT: truncated construct (aa 383-809) after tag removal. GST-HECT 3KR: GST-tagged truncated construct with K-to-R substitution of all three inhibitory lysine residues (aa 383-809; K411R, K432R, K438R). **(C)** *In vitro* ubiquitylation of ^ZIP-MBP^Mga2 at 30°C with different Rsp5 constructs after reconstitution into 100% POPC membranes. Assay components: 70 nM of E1, 500 nM of E2, 200 nM of the indicated Rsp5 construct, 2 µM of ^ZIP-MBP^Mga2, 15 µM of ubiquitin, and ATP. 0.55 µg of ^ZIP-MBP^Mga2 was loaded per lane and quantified by in-gel fluorescence with a Typhoon laser scanner (488 nm laser, Cy2 filter, 25 µm resolution, PMT voltage of 360 V). **(D)** Analysis of construct-dependent Rsp5 autoubiquitylation after in vitro ubiquitylation of ^ZIP-MBP^Mga2. 8.3 ng Rsp5 was loaded per lane and detected by immunoblotting using polyclonal anti-Rsp5 serum and IRDye^®^ 800 CW goat anti-rabbit (1:15000) as the secondary antibody. The blots were scanned on a LI-COR Odyssey^®^ scanner at 700 and 800 nm. Red arrows indicate the unmodified construct at t = 0 min.

To reconstitute the ubiquitylation of ^ZIP-MBP^Mga2, we mixed proteoliposomes with ubiquitin, 1 mM ATP, an ATP-regenerating system, and a set of enzymes for ubiquitin activation (E1), conjugation (E2), and ligation (E3). Due to their known functional equivalence, we could use 6xHistidine-tagged Uba1 from mouse (E1, 70 nM) and human GST-tagged UbcH5B (E2, 500 nM) to activate and deliver ubiquitin to the yeast ubiquitin ligase Rsp5 (E3, 200 nM) (Figure 1E, F). Notably, the E2 identity has no impact on the ubiquitin chains formed by Rsp5 (Kim & Huibregtse, 2009) and this set of proteins was previously used to characterize the mechanism of ubiquitin ligation upon E2-to-E3-to-substrate transfer (Kamadurai *et al*, 2013). We purposely omitted the inclusion of DUBs, such as Ubp2 or Ubp15, which are thought to antagonize Rsp5 and remodel ubiquitin chains after their formation (Kee *et al*, 2005; Ho *et al*, 2017). Different Rsp5 variants (Figure EV1A, B) ubiquitylated ^ZIP-MBP^Mga2 with distinct efficacies (Figure EV1C), while also undergoing autoubiquitylation to a varying degree (Figure EV1D). Compared to full-length Rsp5, a truncated variant (HECT, aa 383-809) lacking the N-terminal C2 domain and the first two of three WW domains (Figure EV1A) provided only a minimal degree of ^ZIP-MBP^Mga2 ubiquitylation (Figure EV1C) in a tightly packed membrane environment (100 mol% POPC), yet it underwent significant autoubiquitylation (Figure EV1D). C2 and WW domains can stabilize non-productive, yet catalytically hyperactive conformations of HECT E3 ligases in the absence of a client protein causing increased E3 ligase autoubiquitylation (Wiesner *et al*, 2007; Wang *et al*, 2010; Mari *et al*, 2014; Riling *et al*, 2015; Zhu *et al*, 2017; Wang *et al*, 2019). We speculate that a more stable engagement between ^ZIP-MBP^Mga2 and full-length Rsp5 promotes ubiquitylation of the lipid saturation sensor (Figure EV1C) whilst lowering Rsp5 autoubiquitylation (Figure EV1D). Notably, the ubiquitylation of Rsp5 at lysine residues K411, K432, or K438 lowers the E3 ligase activity and may serve auto-regulatory purposes (Attali *et al*, 2017). Consistently, the substitution of these three lysines with arginine (3KR) decreases autoubiquitylation of the truncated Rsp5 variant (Figure EV1D) while increasing the ubiquitylation of the lipid saturation sensor (Figure EV1C). For all following experiments, we decided to use wild-type (WT), full-length Rsp5 after removal of the GST-tag, as its presence limited ^ZIP-MBP^Mga2 ubiquitylation (Figure EV1C).

#### Physical and kinetic modeling of Mga2 ubiquitylation

A detailed kinetic analysis of ^ZIP-MBP^Mga2 ubiquitylation requires quantitative data on the abundance of individual ^ZIP-MBP^Mga2-Ub species. To this end, we performed *in vitro* ubiquitylation assays using fluorescently labeled ^ZIP-MBP^Mga2 reconstituted in liposomes composed of tightly packing lipids (100 mol% POPC). We quantified both unmodified and ubiquitylated species by in-gel fluorescence over time (Figure 2A), which neither distinguishes between different linkage types nor between multiple mono- and polyubiquitylations. Because a minimum of 3-6 ubiquitin moieties is required for Cdc48^Npl4-Ufd1^-dependent handling of ubiquitylated substrates and the handover to the proteasome (Thrower *et al*, 2000; Bodnar & Rapoport, 2017; Williams *et al*, 2023; Tsuchiya *et al*, 2017; Kiss *et al*, 2025) and even multi-monoubiquitylations can provide a signal for degradation (Braten *et al*, 2016), we focused our kinetic analysis on ^ZIP-MBP^Mga2-Ub species with up to six ubiquitin.

**Figure 2:**
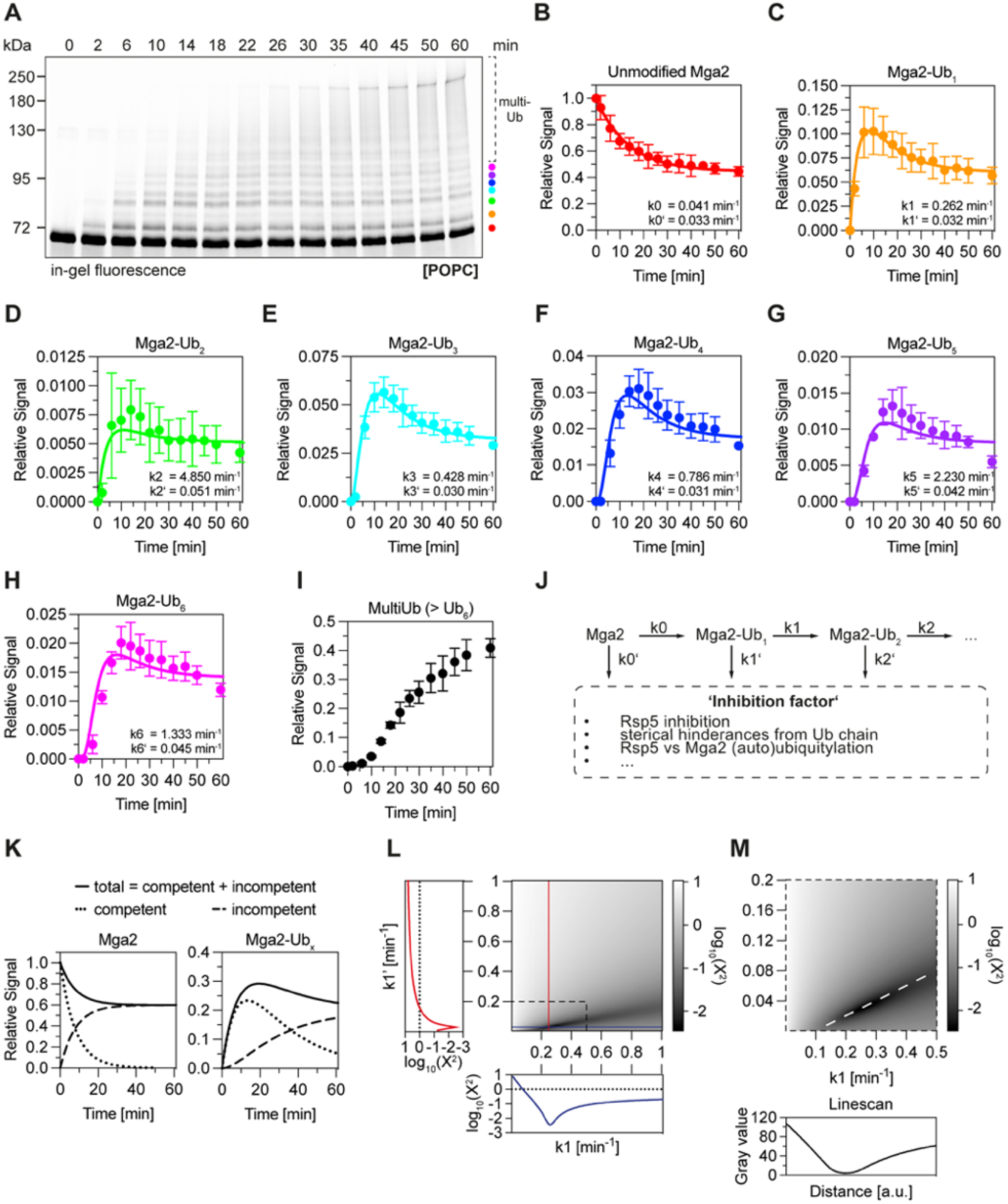
^ZIP-MBP^Mga2 and fitting the ubiquitylation reaction kinetics. **(A)** SDS-PAGE analysis of an *in vitro* ubiquitylation assay with ATTO 488-labeled ^ZIP-MBP^Mga2 in 100 mol% POPC at 30°C. Assay components: 2 µM ^ZIP-MBP^Mga2, 200 nM Rsp5, 500 nM E2 (GST-UbcH5B), 70 nM E1 (6xHis-Uba1), 15 µM ubiquitin (8xHis-Ub), ATP. In total, 0.55 µg of ^ZIP-MBP^Mga2 was loaded for SDS-PAGE. Fluorescence emission was detected with a Typhoon laser scanner (488 nm laser, Cy2 filter, 25 µm resolution, 360 V PMT voltage). Different ^ZIP-MBP^Mga2 species are color-coded. **(B)-(I)** Densitometric quantification of ^ZIP-MBP^Mga2-Ub species over time, normalized to unmodified Mga2 at t = 0 min. The mean and SD of n = 5 replicates is plotted and fitted to the model (solid lines). **(B)** Unmodified ^ZIP-MBP^Mga2, **(C)** ^ZIP-MBP^Mga2-Ub^1^, **(D)** ^ZIP-MBP^Mga2-Ub_2_, **(E)** ^ZIP-MBP^Mga2-Ub_3_, **(F)** ^ZIP-MBP^Mga2-Ub_4_, **(G)** ^ZIP-MBP^Mga2-Ub_5_, **(H)** ^ZIP-MBP^Mga2-Ub_6_, **(I)** multiUb (> Ub_6_) **(J)** Scheme of the kinetic model assuming sequential ubiquitylation with forward rate kx and inhibition rate kx’, with ‘x’ representing the number of ubiquitin modifications on Mga. **(K)** Contribution of ubiquitylation (competent fraction) and inhibition (incompetent fraction) to the fit (total) for unmodified Mga2 and ubiquitylated Mga2-Ub_x_ species. **(L)** Exploration of the parameter space for k1 and k1’ and determination of log_10_(χ^2^) as quality parameter for the goodness of the overall fit. χ^2^ was calculated as the sum of squares of the difference between the mean of the data and the fit, weighted for the standard deviation 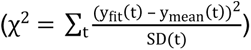. **(M)** Enlarged view of the dashed section shown in **(N)**. White dashed line: line scan across the area containing the optimal solution.

Expectedly, unmodified ^ZIP-MBP^Mga2 was readily consumed over time (Figure 2A, B). However, less than 60% of the protein was modified after 60 min reaction (Figure 2A, B), even though >80% should be accessible to the E3 ligase (Figure 1D). This raised the question, why some ^ZIP-MBP^Mga2 molecules were excluded from ubiquitylation. The abundance of ubiquitylated intermediates (^ZIP-MBP^Mga2-Ub_x_) followed a characteristic time course (Figure 2C-H) with an initial build-up towards a maximal abundance, which was followed by a phase of consumption towards a plateau. Different ^ZIP-MBP^Mga2-Ub species reached their peak abundance consecutively: first, ^ZIP-MBP^Mga2-Ub_1_, then ^ZIP-MBP^Mga2-Ub_2_, then ^ZIP-MBP^Mga2-Ub_3_, and so forth, which is characteristic for HECT domain E3 ligases and consistent with multiple transfers of single ubiquitin molecules (Pierce *et al*, 2009; Deol *et al*, 2019). While multiubiquitylated species with seven or more modifications accumulated over time (Figure 2I), none of the preceding intermediates ^ZIP-MBP^Mga2-Ub_x_ were fully converted to ^ZIP-MBP^Mga2-Ub_x+1_ (Figure 2C-H). Likewise, the formation of highly modified species slowed down at later time points of the reaction (Figure 2I). These observations point toward the existence of one or several inhibitory mechanisms that counteract the complete conversion of reaction intermediates (Figure 2B-H). While neither ATP nor free ubiquitin (Figure EV2A) should be limiting under our experimental conditions, these may be caused by rather technical reasons common to *in vitro* ubiquitylation assays including the inactivation of the E1 or E2 enzymes, or aggregation of the E3 ligase and/or its client. Furthermore, growing ubiquitin chains may limit access to the catalytic site of Rsp5 through steric hindrances and entropic contributions (Brown *et al*, 2015), while different Mga2 molecules with growing ubiquitin chains may compete Rsp5 binding. Last but not least, Rsp5 may undergo an autoubiquitylation that would limit the E3 ligase activity toward its client (Attali *et al*, 2017). In any case, a quantitative description of the reaction kinetics should consider not only the sequential addition of ubiquitin molecules to the lipid saturation sensor, but also inhibitory mechanisms that counteract a complete conversion of reaction intermediates.

To accommodate this requirement, we repurposed the sequential ubiquitylation model for single turnover reactions pioneered by Pierce *et al*. (Pierce *et al*, 2009) and applied it to our steady-state conditions in which the E3 ubiquitin ligase can encounter its client multiple times (Figure 2J). Hence, each reaction intermediate can either receive a new ubiquitin modification with the forward rate kx or undergo an irreversible inhibition with the rate kx^’^, which serves as a collective term for a variety of possible inhibitory mechanisms. It follows that each ubiquitylated ^ZIP-MBP^Mga2 intermediate exists in two pools: one that is ubiquitylation-competent, and one that is inhibited and therefore excluded from further rounds of ubiquitylation (incompetent) (Figure 2K).

The model robustly recapitulates the abundance of ubiquitylation intermediates over time with a characteristic initial accumulation of ^ZIP-MBP^Mga2-Ub_x_ towards a peak abundance, and a subsequent consumption towards a plateau (Figure 2B-H). Global fitting of our data using the average abundance of each intermediate at each time point (Figure 2B-H) was performed to estimate the individual rates for forward ubiquitylation and irreversible inhibition. We found that the first addition of ubiquitin is particularly slow (k0 = 0.041 min^-1^) and that the rate of ubiquitin transfer increases dramatically at later steps yielding 6.4-fold (k1 = 0.262 min^-1^) and even 120-fold higher rates (k2 = 4.85 min^-1^) for ubiquitin transfer. These data suggest that the addition of the first ubiquitin is rate-limiting and key to support subsequent rounds of ubiquitylation, e.g. by increasing the local density of ubiquitin attachment sites for the rather unselective E3 ubiquitin ligase Rsp5, which can generate multiple ubiquitin linkage types (Saeki *et al*, 2009; Kim & Huibregtse, 2009; French *et al*, 2009; Fang *et al*, 2016; Sardana *et al*, 2019). In contrast to the impressive acceleration of the forward reaction, the inhibitory rates k0’ to k6’ were nearly identical within a factor of 2 for all reaction steps (≈0.03 min^-1^ to 0.05 min^-1^) and in the same order of magnitude as the rate-limiting step of ubiquitylation (0.041 min^-1^). Hence, the inhibitory mechanism seems to be insensitive to the number of ubiquitins attached to the sensor. The relevance of the inhibitory rates for the shape of curves in our time course experiments (Figure 2B-H) becomes more obvious when the ubiquitylation-competent pool of ^ZIP-MBP^Mga2 and the inhibited pool are plotted individually (Figure 2K; Supplementary Figure S2A-G): without any inhibitory mechanisms at work, each reaction intermediate of ^ZIP-MBP^Mga2-Ub_x_ would be fully consumed rather than approaching a steady-state concentration. Hence, the inclusion of an inhibitory mechanism is essential to reliably fit our experimental data. More importantly, the combination of an accelerating forward reaction with an inhibitory feedback mechanism fulfills an important prerequisite for signal amplification based on the ubiquitylation kinetics (Ferrell & Xiong, 2001).

We wanted to test the validity of our 14-parameter fit and learn if the extracted rates indeed represent an optimal solution. To this end, we established a quality parameter, log_10_(χ^2^), which is calculated as the sum of squared deviations of the fit to the average abundance at each time point, weighted by the standard deviation 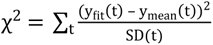. While the rates k0 and k0’ can be analytically established, the rates of the following steps are affected by all preceding rates that determine how fast a certain reaction intermediate is formed and ‘replenished’. We decided to test the quality of the fit progressively by recapitulating the ubiquitylation reaction step by step. Initially, we fixed the analytically determined rates k0 and k0’ to systematically explore a wide parameter space for the forward and inhibition rates k1 and k1’, respectively, while plotting the resulting log_10_(χ^2^) after global fitting (Figure 2L). This analysis revealed a clearly defined minimum of the quality parameter log_10_(χ^2^) for both k1 and k1’ indicating optimal solutions (Figure 2L). A possible caveat of this analysis, however, is that the rates k1 and k1’ could degenerate: an overestimation of the forward rate k1 may be compensated by an equivalent overestimation of the inhibitory rate k1’, such that good fits are achieved whenever a certain ratio of k1 to k1’ is preserved. Indeed, systematically plotting the quality parameter log_10_(χ^2^) for k1 and k1’ pairs identified a strong correlation between k1 and k1’ for yielding good fits (Figure 2L). Nonetheless, even along the diagonal of k1 and k1’ pairs yielding good fits, we found a defined minimum of log_10_(χ^2^) (Figure 2M), hence indicating that the estimated rates are indeed derived from an optimal solution. We then fixed all rates up until this step and systematically explored the parameter space for k2 and k2’, then for k3 and k3’ and so forth to consecutively identify their impact on the overall quality of the fit (Supplementary Figure S2H-L). This analysis demonstrated that the rates derived from the best solution are particularly reliable for the first five rounds of ubiquitylation. In summary, our detailed kinetic analysis reveals a remarkable acceleration of forward ubiquitylation and uncovers an important contribution of an irreversible inhibition for describing the reaction. Notably, the overall fidelity of the reaction was greatly affected when a lysine-less variant of ubiquitin was used instead of WT ubiquitin (Figure EV2B, C). The abundance of high molecular species of ^ZIP-MBP^Mga2 reaction was greatly reduced (Figure EV2C), while the rate of the first ubiquitin attachment was indistinguishable for both ubiquitin variants (Figure EV2D). Later intermediates of the reaction showed vastly distinct reaction kinetics (Figure EV2E-J) leading ultimately to significantly lower levels of high molecular weight ^ZIP-MBP^Mga2 modified with seven or more ubiquitins (Figure EV2K).

Even though we were unable to deduce the exact rates at each ubiquitylation step due to parameter degeneracy, our data show that lysine-less ubiquitin has a major impact on the reaction. Furthermore, these findings suggest that polyubiquitin chains contribute significantly to the pool of highly modified ^ZIP-MBP^Mga2 even though our in-gel fluorescence readout does not distinguish between multi-mono- and polyubiquitylation.

**Figure EV2:**
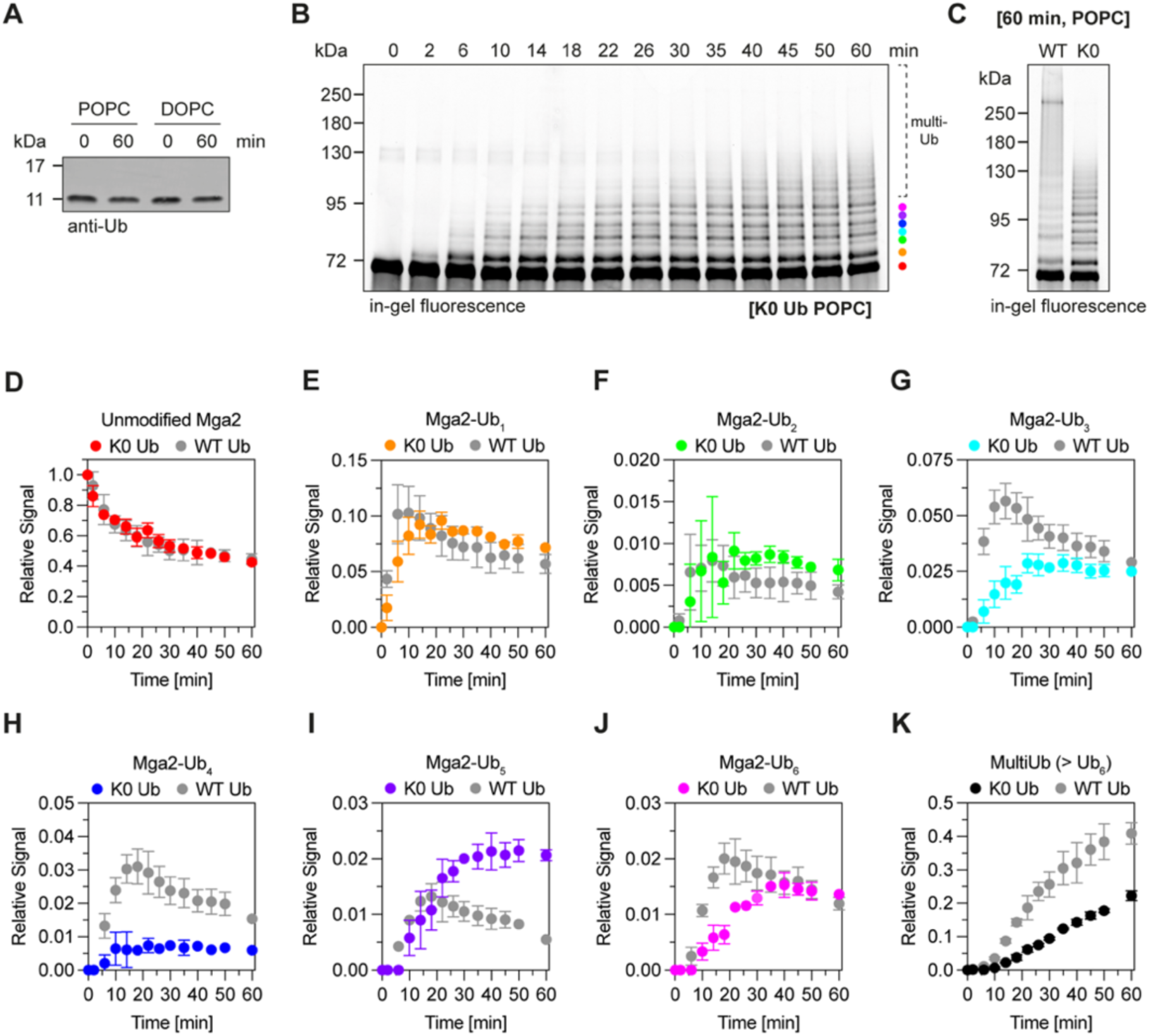
Ubiquitylation of ^ZIP-MBP^Mga2 using lysine-less ubiquitin. **(A)** Detection of free ubiquitin after *in vitro* ubiquitylation of ^ZIP-MBP^Mga2. Samples from Fig. 2A (POPC) and 3C (DOPC) containing 89 ng of ubiquitin were analyzed by SDS-PAGE and immunoblotting. Primary antibody: anti-ubiquitin (1:1000, rabbit). Secondary antibody: IRDye 800 CW^®^ anti-rabbit (1:15,000, goat). **(B)** SDS-PAGE analysis of ubiquitylation ^ZIP-MBP^Mga2 reconstituted into POPC members using lysine-less (K0) ubiquitin. Assay components: 2 µM ^ZIP-MBP^Mga2, 200 nM Rsp5, 500 nM E2, 70 nM E1, 15 µM K0 ubiquitin (8xHis-tagged), ATP. Reactions were incubated at 30°C. Per sample, 0.55 µg of ^ZIP-MBP^Mga2 were loaded for SDS-PAGE (gel: 7.5% Mini-PROTEAN^®^ TGX™). Fluorescent ^ZIP-MBP^Mga2 species were detected using a Typhoon laser scanner (488 nm laser, Cy2 filter, 25 µm resolution, 360 V PMT voltage). **(C)** Comparison of ^ZIP-MBP^Mga2 ubiquitylation using WT and lysine-less (K0) ubiquitin after 60 min. **(D)-(K)** Quantification of ^ZIP-MBP^Mga2 ubiquitylation. Fluorescence intensity was normalized to the signal of unmodified Mga2 at time point t = 0 min. Shown are the mean and SD of n = 3 independent experiments. Data from ubiquitylation assays performed with WT ubiquitin are included for comparison (gray). **(D)** Unmodified ^ZIP-MBP^Mga2, **(E)** ^ZIP-MBP^Mga2-Ub_1_, **(F)** ^ZIP-MBP^Mga2-Ub_2_, **(G)** ^ZIP-MBP^Mga2-Ub_3_, **(H)** ^ZIP-MBP^Mga2-Ub_4_, **(I)** ^ZIP-MBP^Mga2-Ub_5_, **(J)** ^ZIP-MBP^Mga2-Ub_6_, **(K)** multiUb (> Ub_6_).

### Distinguishing lipid environments through Mga2 ubiquitylation

We wanted to assess if lipid saturation affects the kinetics of ^ZIP-MBP^Mga2 ubiquitylation. Previously, it was shown that the supplementation with UFAs to the culture medium abolishes Mga2 processing *in vivo* (Jiang *et al*, 2002; Covino *et al*, 2016) and that the *in vitro* ubiquitylation of ^ZIP-MBP^Mga2 is controlled by lipid packing (Ballweg *et al*, 2020) (Figure 3A). We confirmed a significant impact of lipid packing on ^ZIP-MBP^Mga2 ubiquitylation in our fully defined *in vitro* system (Figure 3B).

**Figure 3:**
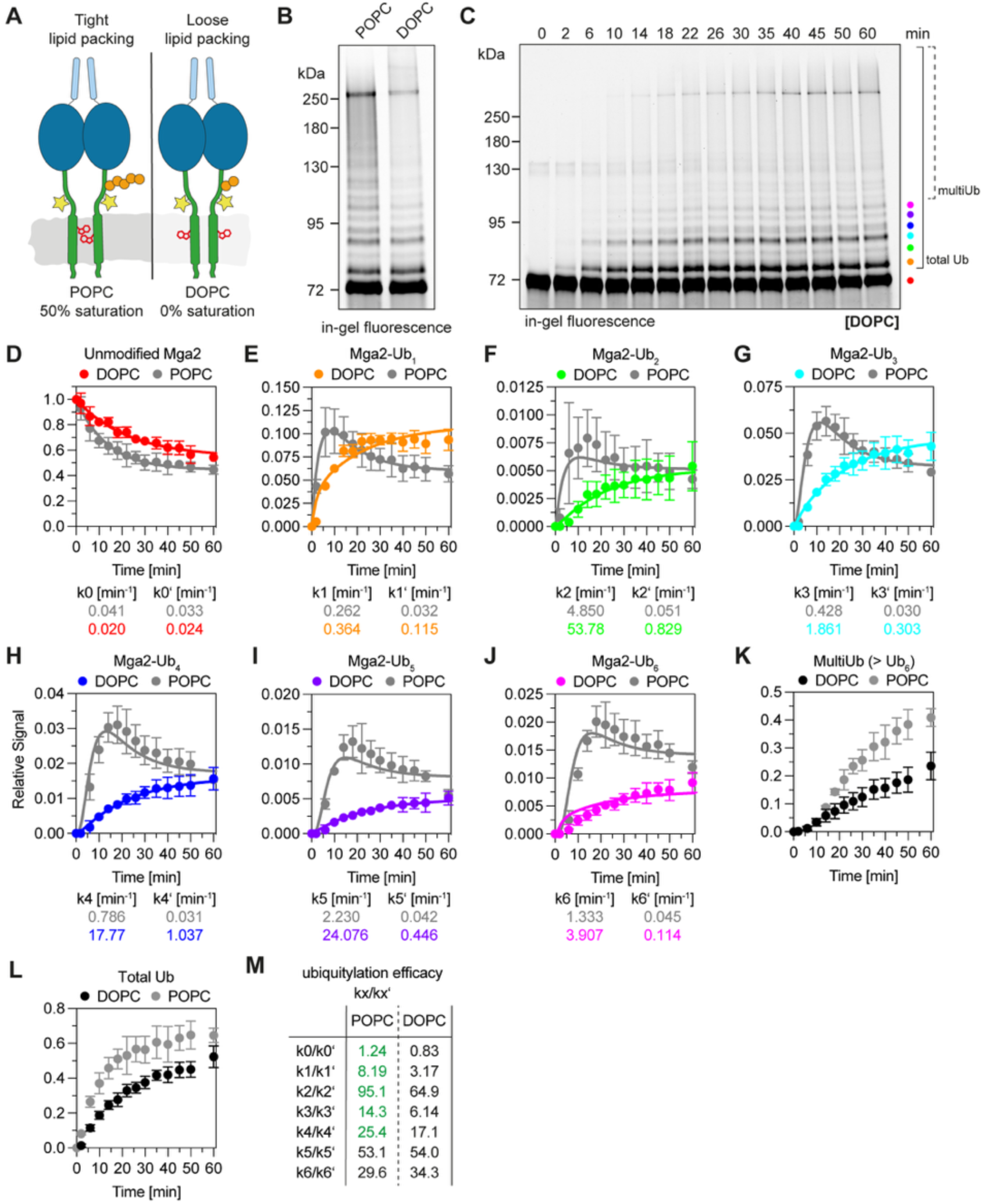
Reconstituting and kinetic modeling of lipid packing-dependent Mga2 ubiquitylation. **(A)** Schematic of lipid packing sensing by Mga2 using a sensory tryptophan residue in the hydrophobic core of the membrane. *Left*: In densely packed, saturated membranes (POPC), the sensory tryptophan is more likely to ‘hide’ from the lipid environment in the dimer interface, and Mga2 can be ubiquitylated. *Right*: In loosely packed, unsaturated membranes (DOPC), the sensory tryptophan residues spend more time facing the loosely packed lipid environment, thereby lowering the efficacy of Mga2 ubiquitylation. **(B)** Comparison of ^ZIP-MBP^Mga2 ubiquitylation in tightly packed POPC versus loosely packed DOPC membranes after 60 min of ubiquitylation. Assay components: 2 µM ^ZIP-MBP^Mga2, 200 nM Rsp5, 500 nM E2 (GST-UbcH5B), 70 nM E1 (6xHis-Uba1), 15 µM ubiquitin (8xHis-Ub), ATP. Of each sample, 0.55 µg of ^ZIP-MBP^Mga2 in 6.0 µL were analyzed by SDS-PAGE and detected using a Typhoon laser scanner (488 nm laser, Cy2 filter, 25 µm resolution, 360 V PMT voltage). **(C)** *In vitro* ubiquitylation of ^ZIP-MBP^Mga2 in 100 mol% DOPC membranes. Assay components as described in (B). Analysis of 0.55 µg of ^ZIP-MBP^Mga2 by SDS-PAGE (gel: 7.5% Criterion™ TGX™) followed by in-gel fluorescence detection using a Typhoon laser scanner (488 nm laser, Cy2 filter, 25 µm resolution, 360 V PMT voltage). Different ^ZIP-MBP^Mga2 species are color-coded. **(D)-(L)** Densitometric measurement of ^ZIP-MBP^Mga2 species fluorescence, normalized to unmodified ^ZIP-MBP^Mga2 at t = 0 min. Mean and SD of n = 5 experiments. ^ZIP-MBP^Mga2 species are color-coded as in (B). Comparison with ubiquitylation in POPC membranes (in gray, from Figure 2). Data fitted to the theoretical model (solid lines) with optimal rates indicated. **(D)** Unmodified ^ZIP-MBP^Mga2, **(E)** ^ZIP-MBP^Mga2-Ub_1_, **(F)** ^ZIP-MBP^Mga2-Ub_2_, **(G)** ^ZIP-MBP^Mga2-Ub_3_, **(H)** ^ZIP-MBP^Mga2-Ub_4_, **(I)** ^ZIP-MBP^Mga2-Ub_5_, **(J)** ^ZIP-MBP^Mga2-Ub_6_, **(K)** multiUb (> Ub_6_), **(L)** total ubiquitylated species (≥ Ub_1_). **(M)** Calculation of the kx/kx’ ratio for POPC- and DOPC-based assays. Ratios that are higher with POPC-based membranes are indicated in green.

We wondered how two lipid environments featuring almost identical membrane viscosities can impose such robust changes in Mga2 ubiquitylation (Ballweg *et al*, 2020; Ragaller *et al*, 2024). Hence, we reconstituted ^ZIP-MBP^Mga2 in DOPC-based liposomes, performed *in vitro* time-course ubiquitylation experiments, and quantified ^ZIP-MBP^Mga2-Ub species over time (Figure 3C-L). The consumption of unmodified ^ZIP-MBP^Mga2 was roughly 2-fold slower in the loosely packed DOPC-based membrane environment (k0_DOPC_ = 0.020 min^-1^) compared to the POPC condition (k0_POPC_ = 0.041 min^-1^) (Figure 3D). Intermediates of the ubiquitylation reaction showed robust lipid-dependent differences at the time of their peak abundance (Figure 3E-J) leading ultimately to a 2-fold difference of multiubiquitylated species with more than 6 modifications at the of the reaction (Figure 3K). It is unlikely that the difference of the first, rate-limiting ubiquitylation step is due to different interaction strengths between the E3 ligase and its client, because the lipid environment has no impact on the amount of Rsp5 that can be co-precipitated with ^ZIP-MBP^Mga2 reconstituted in either POPC- and DOPC-based liposomes (Figure EV3A, B). Instead, we hypothesize that lipid-controlled conformational changes affect the accessibility of lysine residues suitable for modification. This would be consistent with lipid-controlled, conformational changes in Mga2 between the LPXY motif recruiting the E3 ligase in one protomer and target lysine residues in the other (Ballweg *et al*, 2020).

Differences in the reaction kinetics in saturated and unsaturated membrane environments became increasingly clear at the following rounds of ubiquitylation (Figure 3E-J). In stark contrast to the characteristic time course observed for each ubiquitylated intermediate in POPC-based membranes with a buildup, peak abundance, consumption, and plateau phase, only a rather slow accumulation of the intermediates toward a plateau was found for the more loosely packed, DOPC-based membrane environment (Figure 3E-J). This suggests major differences in the ubiquitylation kinetics of ^ZIP-MBP^Mga2 imposed by the different lipid environments. Remarkably, the maximal difference in the abundance of ubiquitylated intermediates between the two membrane environments increased with each round of ubiquitylation up to the ^ZIP-MBP^Mga2-Ub_5_ intermediate (Figure 3D-H). This observation suggests a gradual signal amplification over several ubiquitylation steps.

A detailed analysis of the reaction kinetics further underscored this conclusion. Global fitting of the time course data in the loosely packed membrane environment yielded rates for the forward ubiquitylation and the irreversible inhibition at each step of the reaction (Figure 3D-J). We determined the rate k0_DOPC_ = 0.020 min^-1^ for the addition of the first ubiquitin and the equivalent rate of irreversible inhibition k0’_DOPC_ = 0.024 min^-1^. We also found robust evidence for increasing rates of forward ubiquitylation (Figure 3D, E). However, the parameter degeneracy inherent to the kinetics of the system made it impossible to determine both the forward rate kx and the irreversible inhibition kx’ for subsequent reaction steps with certainty. This is because an overestimation of the forward reaction rate kx can be compensated by an equivalent overestimation of the rate of inhibition kx’ (Supplementary Figure S3A-F). To address the challenges arising from this parameter degeneracy, we introduced a dimensionless ubiquitylation efficacy factor kx/kx’, which should cancel out misestimations of the two interdependent rates (Figure 3M, Figure EV3C-I). Comparing the ubiquitylation efficacy factors at each step of the reaction revealed consistently higher values for first five rounds of ubiquitylation in tightly-packed POPC than in loosely-packed DOPC. Hence, each of these steps contributes to signal amplification and the robust distinction between saturated and unsaturated membranes (Figure 3M). Of particular interest is the rate-limiting, first addition of ubiquitin. Here, the ubiquitylation efficacy factor is ≈1.25 for the saturated membrane environment, thus favoring the forward reaction, but only ≈0.75 for the unsaturated membrane, thereby favoring irreversible inhibition. Hence, forward ubiquitylation dominates in the saturated membrane environments, while irreversible inhibition dominates in more loosely packed membranes. Ultimately, these differences contribute to a robust distinction between saturated and unsaturated membranes.

**Figure EV3:**
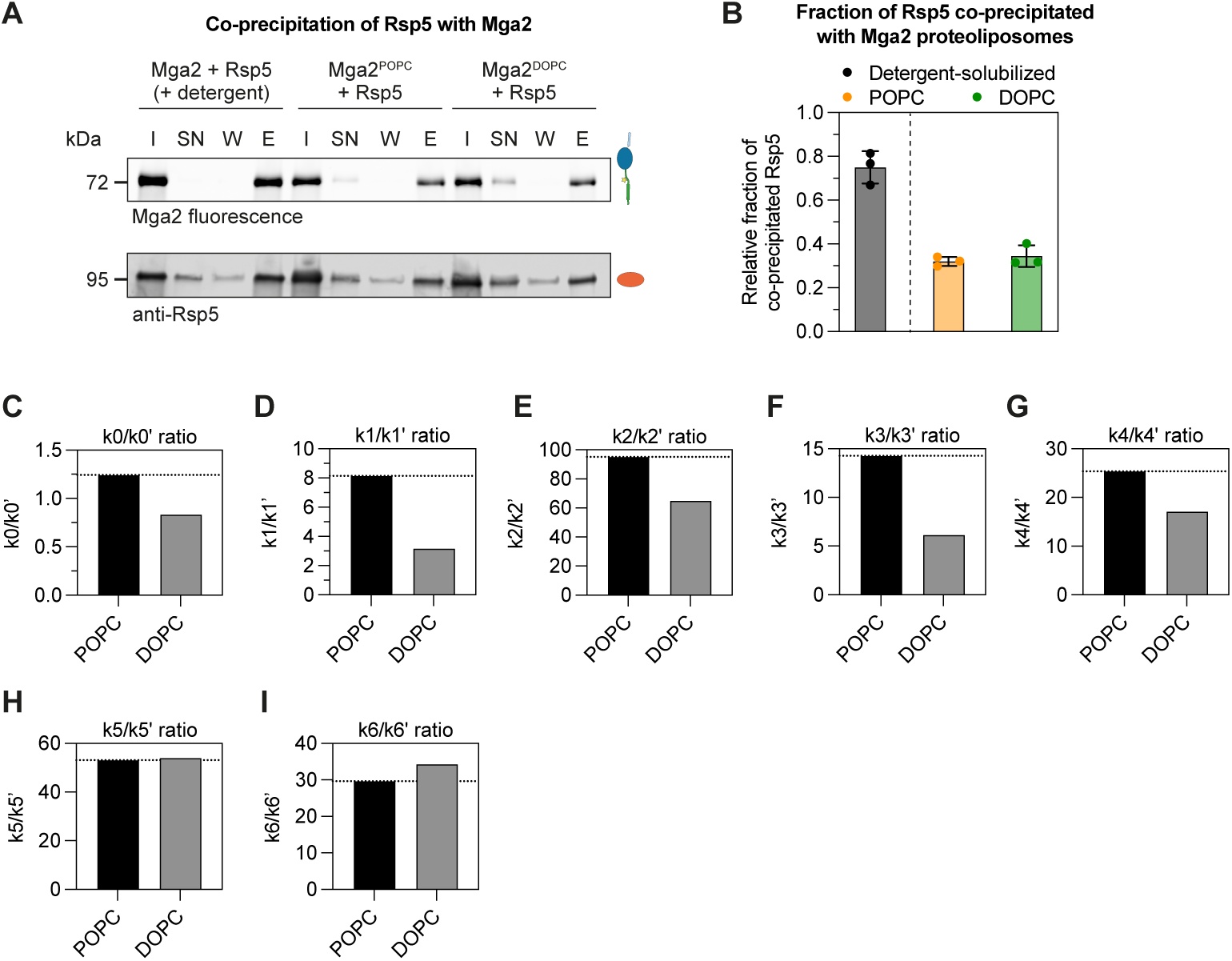
Kinetic distinction between lipid environments and the interaction with Rsp5. **(A)** Co-immunoprecipitation of ^ZIP-MBP^Mga2-containing POPC or DOPC proteoliposomes and Rsp5. Magnetic Dynabeads^®^ were coated with anti-MBP antibody. Input reactions (sample “I”) contained 1 µM ^ZIP-MBP^Mga2 (61.5 µg, in POPC or DOPC proteoliposomes) and 100 nM Rsp5 (9.2 µg). Proteoliposome-containing reactions were compared to a detergent-solubilized condition (50 mM OG). Bound proteins were eluted by adding 1x MSB and 1% SDS, followed by incubation at 95°C for 5 min. SN: supernatant after IP, W: sample after wash step, E: eluate. Fluorescent ^ZIP-MBP^Mga2 species were detected by SDS-PAGE and fluorescence scanning (Typhoon laser scanner, 488 nm laser, Cy2 filter, 360 V PMT, 25 µm resolution). Rsp5 was detected by immunoblotting. Samples containing 83 ng of Rsp5 were analyzed by SDS-PAGE. Primary antibody: anti-Rsp5 serum, 1:1000 (rabbit, polyclonal). Secondary antibody: anti-rabbit IRDye^®^ 800 CW, 1:15000 (goat). Blots were scanned using a LI-COR Odyssey^®^ scanner at 700 and 800 nm (84 µm resolution). **(B)** Quantification of co-immunoprecipitated Rsp5. Signal intensities of eluted Rsp5 (sample “E”) were densitrometrically determined from immunoblots and normalized to input reactions (sample “I”). The mean and SD of n = 3 experiments are shown. **(C) - (I)** Bar graph representation of the kx/kx’ ratio of the optimal rate constants determined for assays with POPC (black) and DOPC (gray) proteoliposomes as provided in Figure 3M.

### The DUB Ubp2 antagonizes Rsp5 but does not affect lipid saturation

In cells, the deubiquitylating enzyme (DUB) Ubp2 is recruited to Rsp5 by the adaptor protein Rup1 and antagonizes Rsp5 by selectively removing K63-linked ubiquitin chains (Kee *et al*, 2005, 2006; Lam *et al*, 2009). In the endocytic pathway, Ubp2 counteracts Rsp5-dependent ubiquitylation of arrestin-related trafficking adaptor proteins together with Ubp15 and Ubp3, with the biggest contribution provided by Ubp2 (Ho *et al*, 2017). Because deubiquitylating activities play crucial roles in decision-making processes such as substrate discrimination in ERAD and substrate ordering by the APC (Rape *et al*, 2006; Zhang *et al*, 2013), we wanted to test if Ubp2 can contribute to lipid saturation sensing by modulating Mga2 ubiquitylation in different lipid environments. We purified recombinant Ubp2 and Rup1 (Figure 4A) and tested their activity in Rsp5 autoubiquitylation assays (Figure 4B,C; Figure EV4A-D). Expectedly, Ubp2 counteracted Rsp5 autoubiquitylation (Figure 4B,C). The antagonistic effect of Ubp2 was titratable (Figure 4B,V) and strongly modulated by Rup1 (Figure 4B,C; Figure EV4A, B), which has no impact on Rsp5 autoubiquitylation on its own (Figure EV4C, D). Hence, Ubp2 is active towards ubiquitylated species of Rsp5.

**Figure 4:**
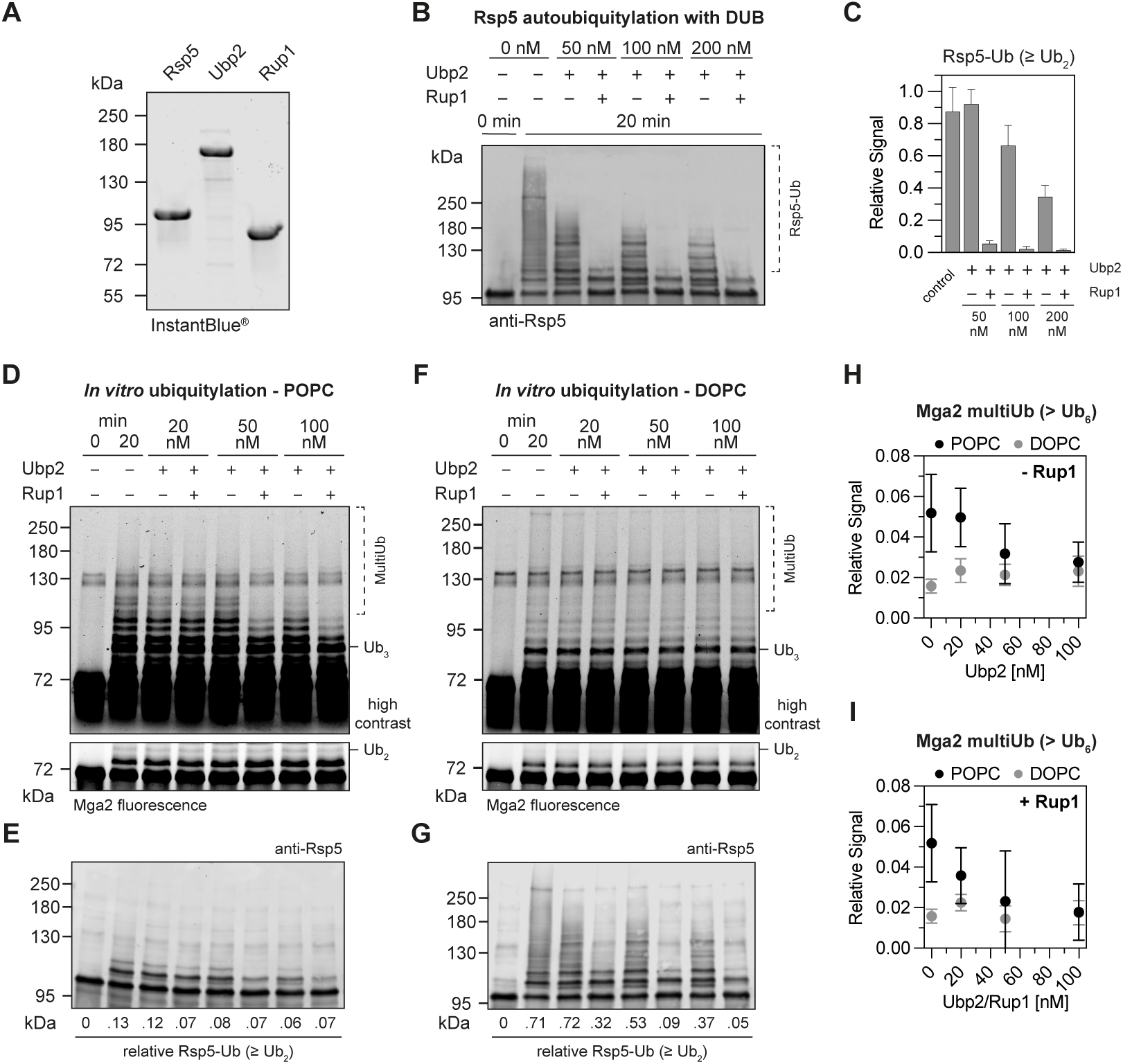
DUB activity reduces saturation-induced differences in Mga2 ubiquitylation. **(A)** Recombinantly produced and purified full-length Rsp5, Ubp2, and Rup1 (0.5 µg/per lane). The gel was stained with IntantBlue^®^. **(B)** *In vitro* Rsp5 autoubiquitylations with Ubp2 and Rup1. 70 nM E1, 500 nM E2, 200 nM Rsp5, 15 µM ubiquitin, and indicated concentrations of Ubp2 and Rup1 were incubated at 30°C for 20 min. 83 ng of Rsp5 were loaded on each lane. The reaction mix was analyzed by SDS-PAGE and immunoblotting using a polyclonal anti-Rsp5 serum (rabbit) an anti-rabbit IRDye 800 CW^®^ (1:15,000, goat) antibody. Blots were scanned on a LI-COR Odyssey^®^ scanner at 700 and 800 nm (84 µm resolution). **(C)** Quantification of Rsp5 autoubiquitylation from assays shown in (A). Background-corrected signals from intermediate molecular weight species (IMW; gray) and high molecular weight species (HMW; black) and normalized to the signal from unmodified Rsp5 at t = 0 min. The mean ± SD of n = 3 experiments are plotted. **(D)** *In vitro* ubiquitylation of ^ZIP-MBP^Mga2 reconstituted in POPC liposomes with Ubp2 and Rup1. 70 nM E1, 500 nM E2, 200 nM Rsp5, 2 µM ^ZIP-MBP^Mga2, 15 µM ubiquitin, and indicated concentrations of Ubp2 and Rup1 were incubated at 30°C for 20 min. 0.55 µg of ^ZIP-MBP^Mga2 were analyzed by SDS-PAGE and in-gel fluorescence scanning (Typhoon laser scanner, 488 nm laser, Cy2 filter, 360 V PMT, 25 µm resolution). **(E)** Samples shown in (D) were used for anti-Rsp5 immunoblotting (8.3 ng of Rsp5 iper lane). The relative fraction of ubiquitylated Rsp5 (≥ Ub_2_) was determined by background-correcting signals using control lanes (t = 0 min), followed by normalization to signals of unmodified Rsp5 at time point t = 0 min. The mean of n = 3 experiments from independent reconstitutions is indicated below each lane. **(F)** *In vitro* ubiquitylation assays ^ZIP-MP^Mga2 reconstituted in DOPC liposomes performed as described in (D). **(G)** Samples from assays shown in (F) containing 8.3 ng of Rsp5 were used for immunoblotting using anti-Rsp5. The relative fraction of ubiquitylated Rsp5 (≥ Ub_2_) was determined as in (C). The mean of n = 3 experiments from independent reconstitutions is provided. **(H)** Quantification of multiUb ^ZIP-MBP^Mga2 species (> Ub_6_) as shown in (D) and (F) from Mga2 ubiquitylation assays in the presence of Ubp2 and with Mga2 reconstituted either in POPC- or DOPC-based liposomes. Background-corrected, normalized fluorescence signals from n = 3 independent reconstitutions are plotted as the mean ± SD. **(I)** Quantification of multiUb ^ZIP-MBP^Mga2 species (> Ub_6_) as shown in (D) and (F) from Mga2 ubiquitylation assays in the presence of indicated concentrations of Ubp2 and Rup1 and with Mga2 reconstituted either in POPC- or DOPC-based liposomes. Background-corrected, normalized fluorescence signals from n = 3 independent experiments are plotted as the mean ± SD.

Next, we tested the impact of Ubp2 and Rup1 on the ubiquitylation of ^ZIP-MBP^Mga2 reconstituted in either POPC-(Figure 4D,E) or in more loosely packed DOPC-based liposomes (Figure 4F,G) (note the shorter assay time of 20 min). With POPC-based proteoliposomes, we observed a mild reduction of ^ZIP-MBP^Mga2 ubiquitylation in the presence of increasing concentrations of Ubp2, which was further reduced by Rup1 (Figure 4D). Likewise, the low degree of Rsp5 autoubiquitylation was further reduced by the presence Ubp2 and Rup1 (Figure 4E). When the same experiments were performed with ^ZIP-MBP^Mga2 reconstituted in DOPC-based liposomes (Figure 4F), we observed less ubiquitylation of ^ZIP-MBP^Mga2 ubiquitylation, but much more Rsp5 autoubiquitylation, which was strongly modulated by the presence of Ubp2 and Rup1 (Figure 4G). Hence, the autoubiquitylation of Rsp5 is affected by the lipid environment of ^ZIP-MBP^Mga2 and can be modulated by the presence of Ubp2 and Rup1.

Quantification of the multiubiquitylated Mga2 species carrying seven or more modifications showed that the largest difference between the two lipid environments was observed in the absence of Ubp2 regardless of the presence of Rup1 (Figure 4H, I). This suggested that Ubp2 and Rup1, while crucially regulating Rsp5 functions *in vivo* (Kee *et al*, 2005; Lam *et al*, 2009; Lam & Emili, 2013; Cavellini *et al*, 2017), play no major role in signal amplification and lipid saturation sensing. This conclusion was validated by *in vivo* experiments (Supplementary Figure S4A-C). Unlike Δ*mga2* or Δ*ole1* cells that showed significant growth defects in synthetic complete dextrose media, Δ*ubp2* and Δ*rup1* cells were indistinguishable from isogenic wildtype cells even when lipid metabolism was perturbed by supplementing the medium with the saturated fatty acid palmitate or unsaturated fatty acid oleate. Consistent with earlier observations, we found that palmitate aggravates the growth defect of Δ*mga2* cells (Surma *et al*, 2013), while oleate restores normal growth of Δ*mga2* cells and rescues the otherwise lethal effect of an Δ*ole1* knockout (Supplementary Figure S4B, C) (Stukey *et al*, 1989; Zhang *et al*, 1999; Hoppe *et al*, 2000; Surma *et al*, 2013; Covino *et al*, 2016). These data imply that Ubp2, even when recruited to Rsp5, does not play major roles in regulating lipid saturation. To test for a more subtle, modulatory role of Ubp2 and Rup1, we characterized wildtype, Δ*ubp2*, and Δ*rup1* cells by quantitative lipidomics (Figure EV4E-H). The lipid compositions of these strains were indistinguishable (Figure EV4E-H) with the only exception that Δ*ubp2* featured mildly increased TAGs for an unknown reason (Figure EV4F) and small change in the length distribution of glycerophospholipid fatty acyl chains (Figure EV4G). Most relevant for the OLE pathway, we found that the degree of lipid saturation was unperturbed in Δ*ubp2* and it Δ*rup1* cells (Figure EV4H). Hence, Ubp2 and Rup1 do not contribute to the regulation of the OLE pathway under standard growth conditions.

**Figure EV4:**
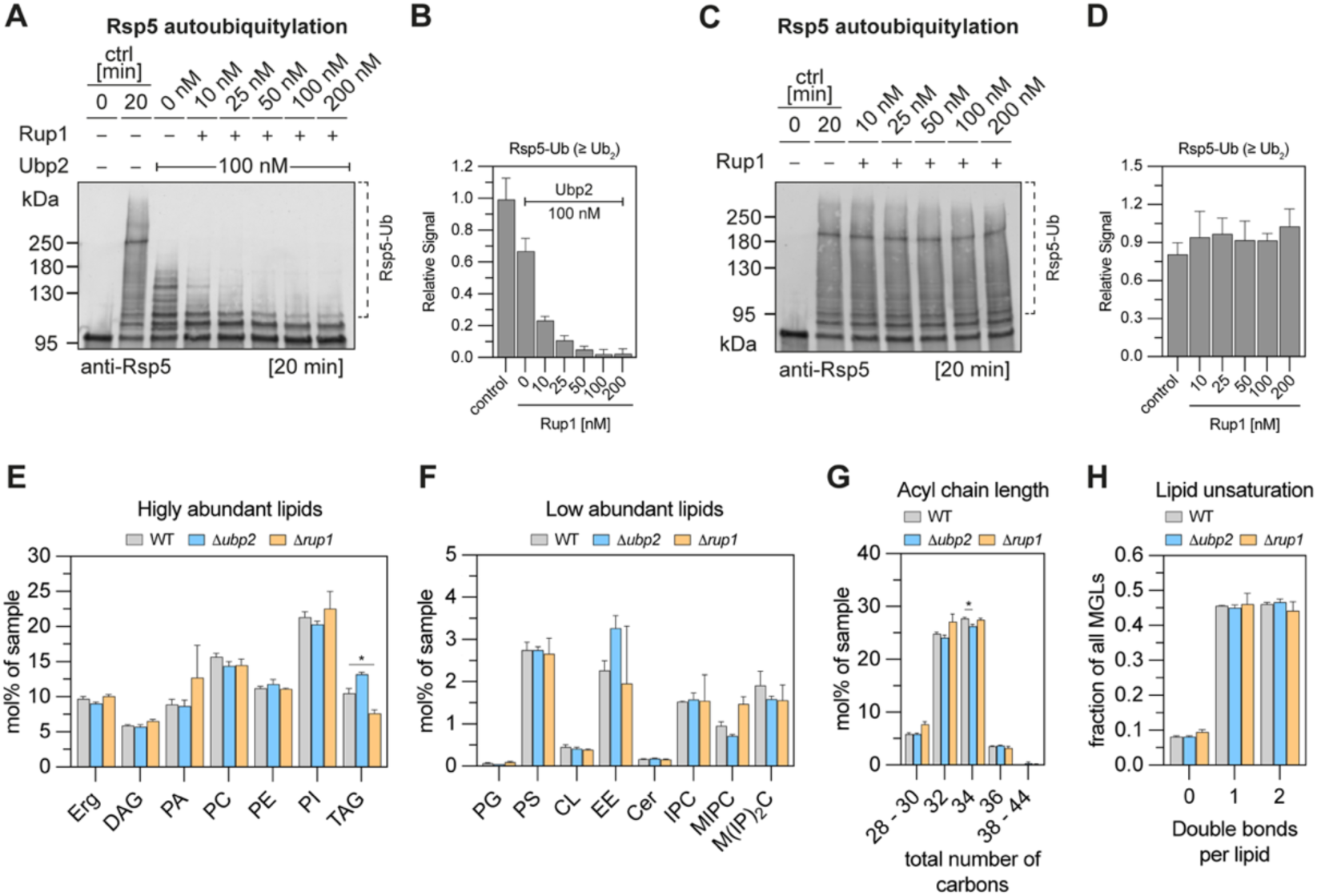
Ubp2 modulates Rsp5 autoubiquitylation with minimal impact on the cellular lipidome. **(A)** *In vitro* autoubiquitylation assay of Rsp5 with Rup1 titration. Assay components: 70 nM E1, 500 nM E2, 200 nM Rsp5, 15 µM ubiquitin, ATP, and indicated concentrations of Ubp2 and Rup1. Assays were performed at 30°C for 20 min. Samples containing 83 ng of Rsp5 were used for SDS-PAGE and immunoblotting. Primary antibody: anti-Rsp5 serum (rabbit, polyclonal). Secondary antibody: anti-rabbit IRDye^®^ 800 CW (goat, 1:15,000). Blots were scanned using a LI-COR Odyssey^®^ scanner at 700 and 800 nm (84 µm resolution). **(B)** Quantification of Rsp5 autoubiquitylation from assays shown in (A). Signals of ubiquitylated Rsp5 (≥ Ub_2_) were background-corrected using control lanes (t = 0 min) and normalized to intensities of unmodified Rsp5 at time point t = 0 min. The mean and SD of n = 3 experiments are shown. **(C)** *In vitro* Rsp5 autoubiquitylation assays with increasing Rup1 concentrations. Assays and subsequent analysis by immunoblotting were performed as described in (F). **(D)** Rsp5 autoubiquitylation (≥ Ub_2_) was determined as described. The mean and SD of n = 3 experiments are shown. **(E) - (H)** Comparing the lipidome of WT yeast and Δ*ubp2* and Δ*rup1* strains. Lipids for lipidomic analyses were isolated from mid-logarithmic yeast cultures (OD_600_ = 0.8). Plotted is the mean and the SD of n = 3 independent experiments. Statistical significances were tested by multiple unpaired t-tests with Welch’s correction using the Holm-Šídák method with a p-value threshold of 0.05. * p < 0.05. **(E)** Relative quantification of highly abundant lipids. Erg: ergosterol, DAG: diacylglycerol, PA: phosphatidic acid, PC: phosphatidylcholine, PE: phosphatidylethanolamine, PI: phosphatidylinositol, TAG: triacylglycerol. For panels (A) to (C), the contribution is given as mol% of all lipids. **(F)** Relative quantification of low abundant lipids. PG: phosphatidylglycerol, PS: phosphatidylserine, CL: cardiolipin, EE: ergosterol esters, CER: ceramide, IPC: inositolphosphorylceramide, MIPC: mannosylinositol-phosphorylceramide, M(IP)_2_C: mannosyl-di-(inositolphosphoryl)ceramide. **(G)** Acyl chain length profile of membrane glycerolipids containing two acyl chains (DAG, PA, PC, PE, PI, PG, and PS). **(H)** Saturation level of membrane glycerolipids (MGLs) containing two acyl chains.

### Ubiquitin at the crossroads – Selective ubiquitylation of either the lipid saturation sensor or Rsp5

Given the central importance of irreversible inhibition for our model of signal amplification, we decided to further characterize Rsp5 autoubiquitylation even though other mechanisms are likely to contribute. As Rsp5 autoubiquitylation appeared sensitive to the lipid environment of Mga2 (Figure 4E, G), we performed time-course experiments with ^ZIP-MBP^Mga2-containing proteoliposomes composed of either tightly packed (POPC) or loosely packed (DOPC) lipids and assayed both ^ZIP-MBP^Mga2 ubiquitylation (Figure 5A) and Rsp5 autoubiquitylation (Figure 5B). Mga2 ubiquitylation was more efficient in the more saturated membrane environment (Figure 5A), while Rsp5 autoubiquitylation proceeded faster in the presence of loosely packed DOPC-based proteoliposomes (Figure 5B). A densitometric analysis, which is only semi-quantitative for high molecular weight smear, confirmed that unmodified Rsp5 was consumed much faster in the presence of DOPC-based proteoliposomes (Figure 5C) resulting in different levels of highly ubiquitylated Rsp5 (Figure 5D).

**Figure 5:**
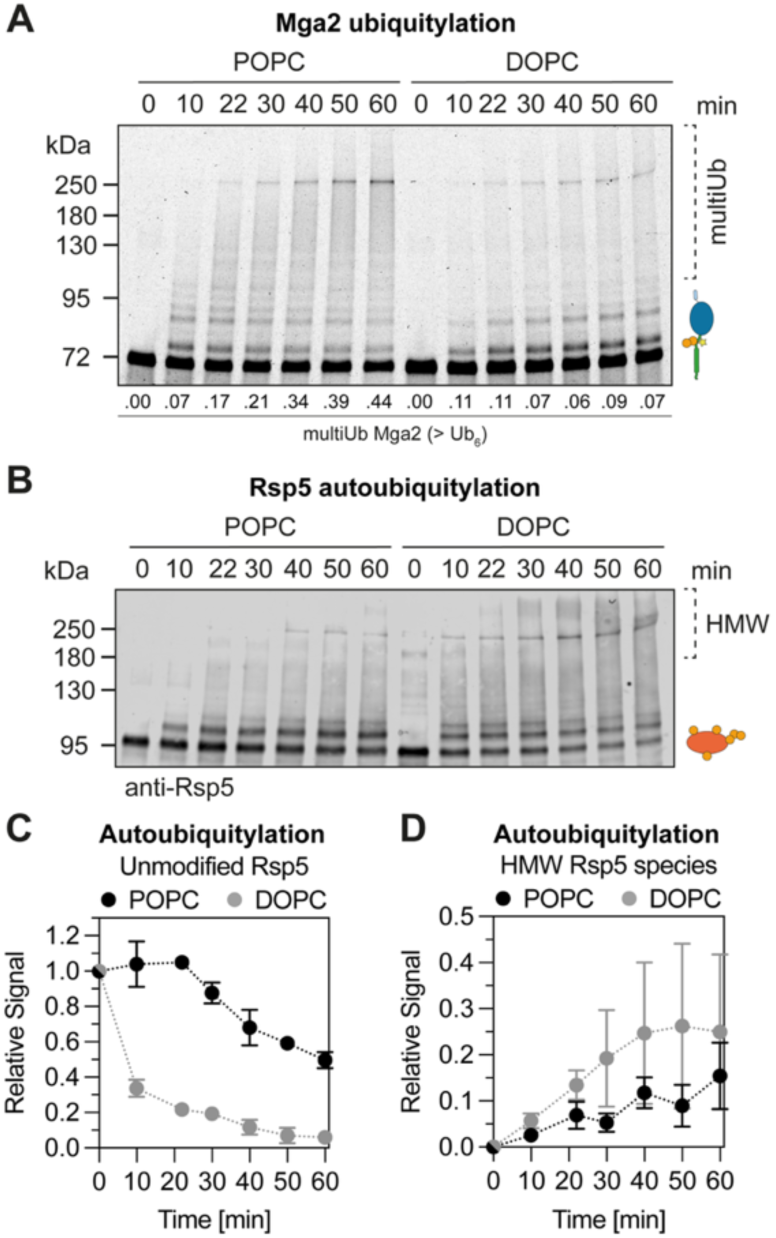
Lipid packing redirects ubiquitylation. **(A)** Comparing the ubiquitylation of ^ZIP-MBP^Mga2 in two different lipid environments. ^ZIP-MBP^Mga2 was reconstituted into tightly packing POPC or loosely packed DOPC membranes. Assay components: 2 µM ^ZIP-MBP^Mga2 (in POPC or DOPC), 200 nM Rsp5, 500 nM E2, 70 nM E1, 15 µM ubiquitin, ATP. Per time point, 0.55 µg of ^ZIP-MBP^Mga2 were loaded for SDS-PAGE (gels: 7.5% Mini-PROTEAN^®^ TGX™). Fluorescent ^ZIP-MBP^Mga2 species were detected using a Typhoon laser scanner (488 nm laser, Cy2 filter, 25 µm resolution, PMT voltage of 360 V). **(B)** For detection of Rsp5, the samples of the *in vitro* ubiquitylation assay were diluted 1:10 and 8.3 ng of Rsp5 were used for SDS-PAGE and anti-Rsp5 immunoblotting. Primary antibody: anti-Rsp5 (rabbit, polyclonal serum 1:1000). Secondary antibody: IRDye^®^ 800 CW goat anti-rabbit (1:15000). The blots were scanned using a LI-COR Odyssey^®^ scanner at 700 and 800 nm (84 µm resolution). **(C)** Quantifying the relative abundance of unmodified Rsp5 from blots shown in (B). Signals were normalized to the intensity of unmodified Rsp5 at time point t = 0 min. Shown are the mean and SD of n = 3 experiments. **(D)** Quantification of high molecular weight (HMW) multiUb Rsp5 species (MW > 180 kDa).

**Figure EV5:**
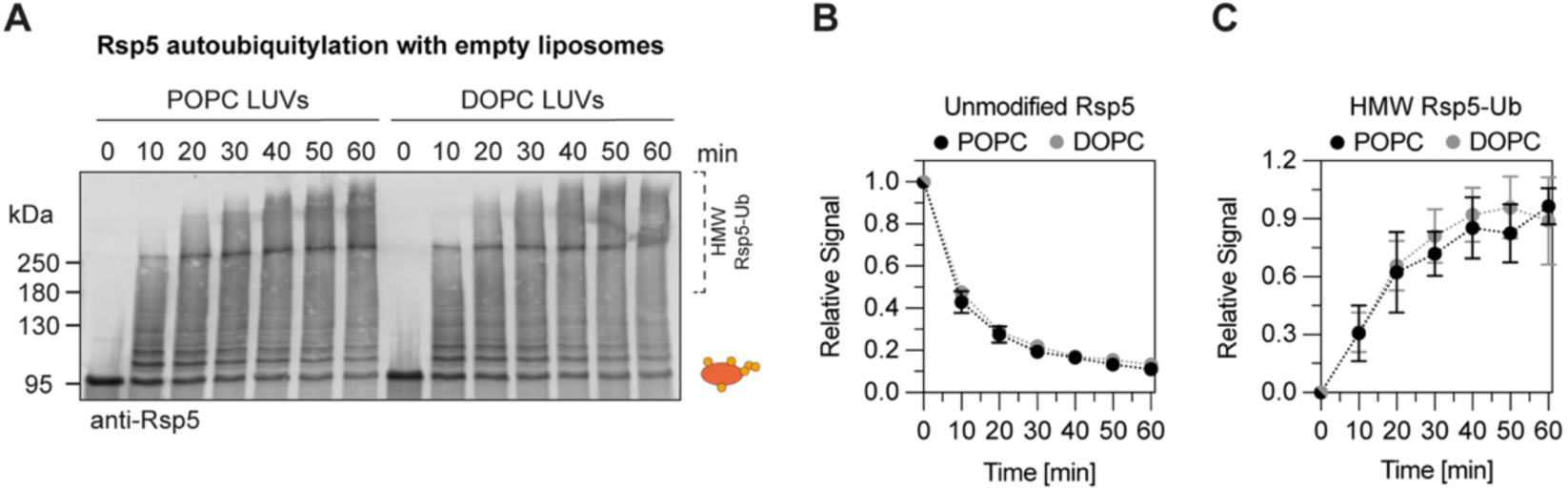
Rsp5 is insensitive to lipid saturation. **(A)** Rsp5 autoubiquitylation assays in the presence of empty large unilamellar vesicles (LUVs) consisting of POPC or DOPC. Assay components: 200 nM Rsp5, 500 nM E2, 70 nM E1, 15 µM ubiquitin, ATP, 16 mM lipids. Lipids were prepared from multilamellar vesicles by extrusion (100 nm filter). Assays were performed at 30°C. 83 ng of Rsp5 were used for SDS-PAGE (gels: 7.5% Mini-PROTEAN^®^ TGX™), followed by immunoblot detection. Primary antibody: anti-Rsp5 serum, 1:1000 (rabbit, polyclonal). Secondary antibody: anti-rabbit IRDye^®^ 800 CW, 1:15000 (goat). Blots were scanned using a LI-COR Odyssey^®^ scanner at 700 and 800 nm (84 µm resolution). **(B)** Quantification of unmodified Rsp5 from blots shown in (A). Signal intensities were normalized to signals of unmodified Rsp5 at time point t = 0 min. The mean and SD of n = 3 experiments are shown. **(C)** Quantification of ubiquitylated Rsp5 from assays shown in (A). Signals of Rsp5-Ub (≥ Ub_2_) were background-corrected using signals from control lanes (t = 0 min) and normalized to the intensity of unmodified Rsp5 at time point t = 0 min. The mean and SD of n = 3 experiments are shown.

Rsp5 contains a C2 domain, which has been implicated in lipid binding and regulating endocytosis by directing client protein ubiquitylation (Dunn & Hicke, 2001; Dunn *et al*, 2004). To test if Rsp5 may sense the lipid packing directly, we performed autoubiquitylation assays in the presence of extruded, protein-free liposomes composed of either POPC or DOPC (Figure EV5A-C). No differences in Rsp5 autoubiquitylation were observed under these conditions (Figure EV5A-C). Hence, the lipid saturation sensor Mga2 is crucial for the lipid-sensitive autoubiquitylation of Rsp5.

Our findings suggest that ubiquitin, that is delivered to the active site of Rsp5, can either be used for ^ZIP-MBP^Mga2 ubiquitylation or Rsp5 autoubiquitylation. The relative efficacy of these processes is dependent on the lipid environment of Mga2 by modulating the structural dynamics of the lipid saturation sensor (Covino *et al*, 2016; Ballweg *et al*, 2020). It is unlikely that the exposure of the LPKY motif in Mga2 is key to the observed differences, as Rsp5 interacts equally well with Mga2 irrespective of the lipid environment (Figure EV3A,B). Instead, our findings suggest that Rsp5 autoubiquitylation is controlled by how effectively Mga2 ‘accepts’ ubiquitin modifications.

### Detailed Characterization of Mga2 and Rsp5 ubiquitylation by mass spectrometry

To further dissect the ubiquitylation of ^ZIP-MBP^Mga2 and Rsp5, we performed *in vitro* ubiquitylation assays for 20 min with ^ZIP-MBP^Mga2 in either POPC- or DOPC-based membrane environments and subjected the reaction mix to mass spectrometry-based analyses (Figure EV6A, B). This revealed the formation of di-ubiquitin linkages via K11, K48, K63, and less so K27 (Figure EV6A, B). These observations are consistent with previous reports (Kim & Huibregtse, 2009; Fang *et al*, 2016; French *et al*, 2009) and indicative for an Rsp5-dependent formation of mixed ubiquitin chains. Furthermore, they suggest a preference for chain formation over multi-monoubiquitylation (Figure EV6C). This interpretation is further supported by results from Rsp5 autoubiquitylation using either wild type (WT), single lysine (K6, K11, K27, K33, K48, K63), lysine-less (K0) or a methylated ubiquitin in the assay, in which autoubiquitylation was most effective with WT ubiquitin, less efficient with single-lysine variants, and most ineffective lysine-less or even a methylated ubiquitin variant (Figure EV6C). It seems that the access to unmodified lysine residues determines the effectiveness of ubiquitylation by Rsp5.

Using mass spectrometry, we also characterized which lysine residues in ^ZIP-MBP^Mga2 are modified during the reaction and studied the impact of the lipid environment (Figure 6A; EV6D, E). Not surprising for an *in vitro* assay and for a client protein with 50 lysine residues, we observed modifications across the entire fusion protein including the N-terminal leucine zipper, the MBP tag, and Mga2 (Figure 6A; EV6D, E). Of these, the residues K980 and K983 from Mga2 were particularly interesting, as they were previously identified as primary sites of ubiquitylation *in vivo* (Bhattacharya *et al*, 2009) (Figure 6A). K980 and K983 modifications were elevated approximately 1.5-fold in the more saturated membrane compared to the unsaturated one (Figure 6A; EV6D), which is consistent with our quantifications by in-gel fluorescence after 20 min *in vitro* ubiquitylation (Figure 3K, L). Similar increases were also observed for non-native lysines such as K339 in the MBP tag (Figure 6A; EV6D). Overall, these results underscore that membrane saturation promotes ^ZIP-MBP^Mga2 ubiquitylation.

**Figure 6:**
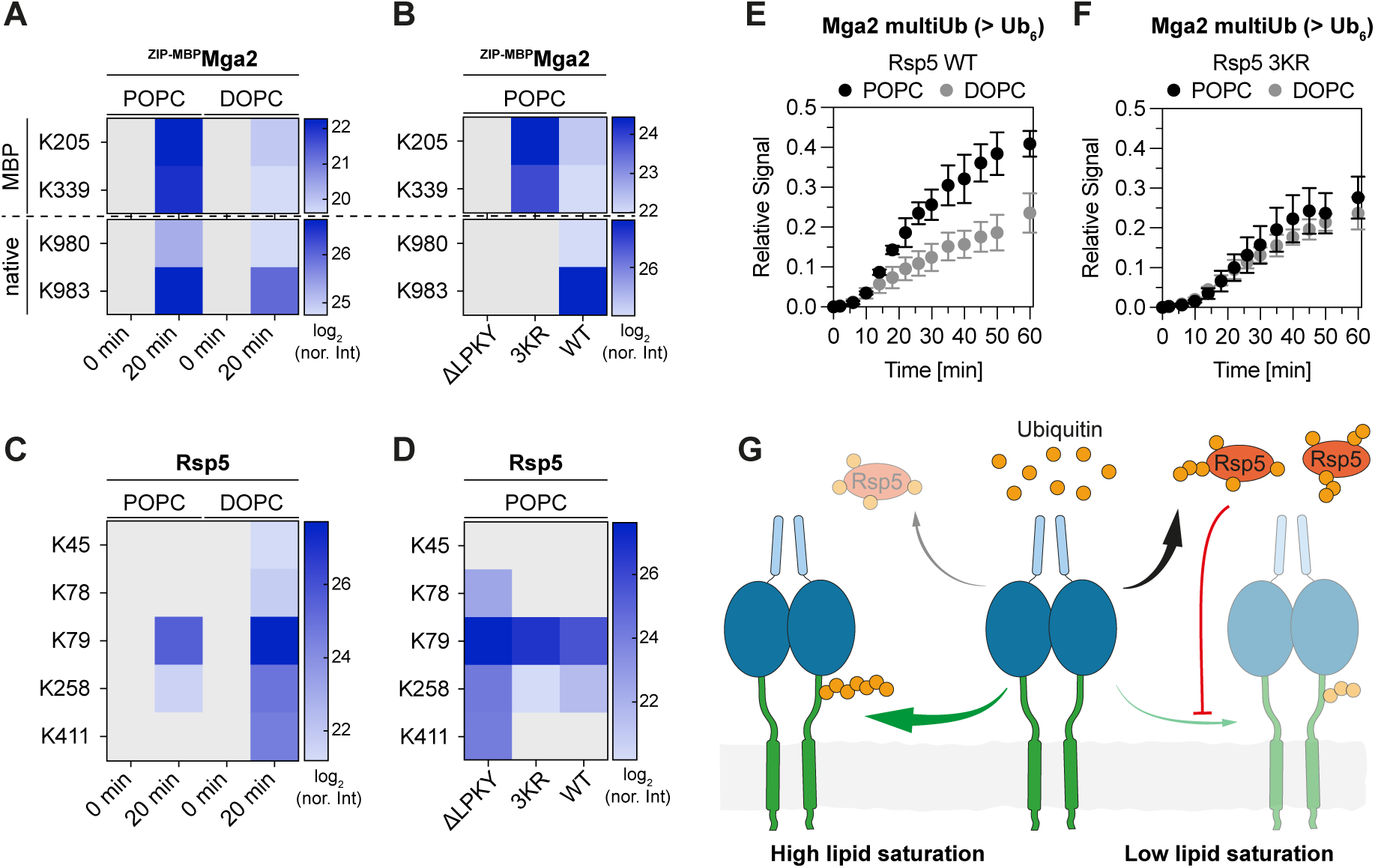
Rsp5 auto-inhibition drives signal amplification during lipid saturation sensing. **(A)** Mass spectrometry analysis of membrane-dependent ^ZIP-MBP^Mga2 ubiquitylation. ^ZIP-MBP^Mga2 was reconstituted into POPC and DOPC membranes. *In vitro* ubiquitylation assays contained 70 nM E1, 500 nM E2, 200 nM Rsp5, 2 µM ^ZIP-MBP^Mga2 (in DOPC or POPC), 15 µM ubiquitin, and ATP. Reactions were incubated at 30°C. The signal was normalized to the total intensity. Shown is the log_2_ of the normalized intensities of n = 3 replicates. **(B)** Mass spectrometric analysis of ^ZIP-MBP^Mga2 ubiquitylation using mutants. Assays were performed as detailed in (A), using POPC-reconstituted WT or mutant ^ZIP-MBP^Mga2 constructs. ΔLPKY: deletion of the Rsp5 binding motif from Mga2. 3KR: triple lysine-to-arginine substitution of Mga2’s native lysines (K980R, K983R, K985R). **(C) - (D)** Mass spectrometric analysis of Rsp5 autoubiquitylation from assays shown in (A) and (B). **(E)** Analysis of membrane-dependent ^ZIP-MBP^Mga2 multiubiquitylation (> Ub_6_) using WT Rsp5 and ^ZIP-MBP^Mga2 reconstituted into POPC and DOPC membranes. Data are reproduced from Fig. 3. **(F)** *In vitro* ubiquitylation of ^ZIP-MBP^Mga2 using an Rsp5 triple arginine-to-lysine mutant blocking autoubiquitylation-induced self-inhibition. Assay components: 70 nM E1, 500 nM E2, 200 nM Rsp5 3KR (K411R, K432R, K438R), 2 µM ^ZIP-MBP^Mga2 (in POPC or DOPC), 15 µM ubiquitin, ATP. Reactions were incubated at 30°C. Samples were analyzed by SDS-PAGE and in-gel fluorescence scanning. Signal intensities were normalized to the signal of unmodified ^ZIP-MBP^Mga2 at time point t = 0 min. The mean and SD of n = 3 experiments are shown. **(G)** Schematic model of signal amplification during lipid saturation sensing by Mga2. Left: Saturated, tightly packing membranes support the ubiquitylation of Mga2. The accelerating reaction rates create positive feedback that sustains amplification of membrane-derived signals. Right: Ubiquitylation of Mga2 is dominated by inhibition when lipid saturation and packing are low, leading to the pronounced autoubiquitylation of Rsp5. Ligase self-modification inhibits and competes with Mga2 ubiquitylation, ultimately resulting in the reduction of Mga2 modification.

**Figure EV6:**
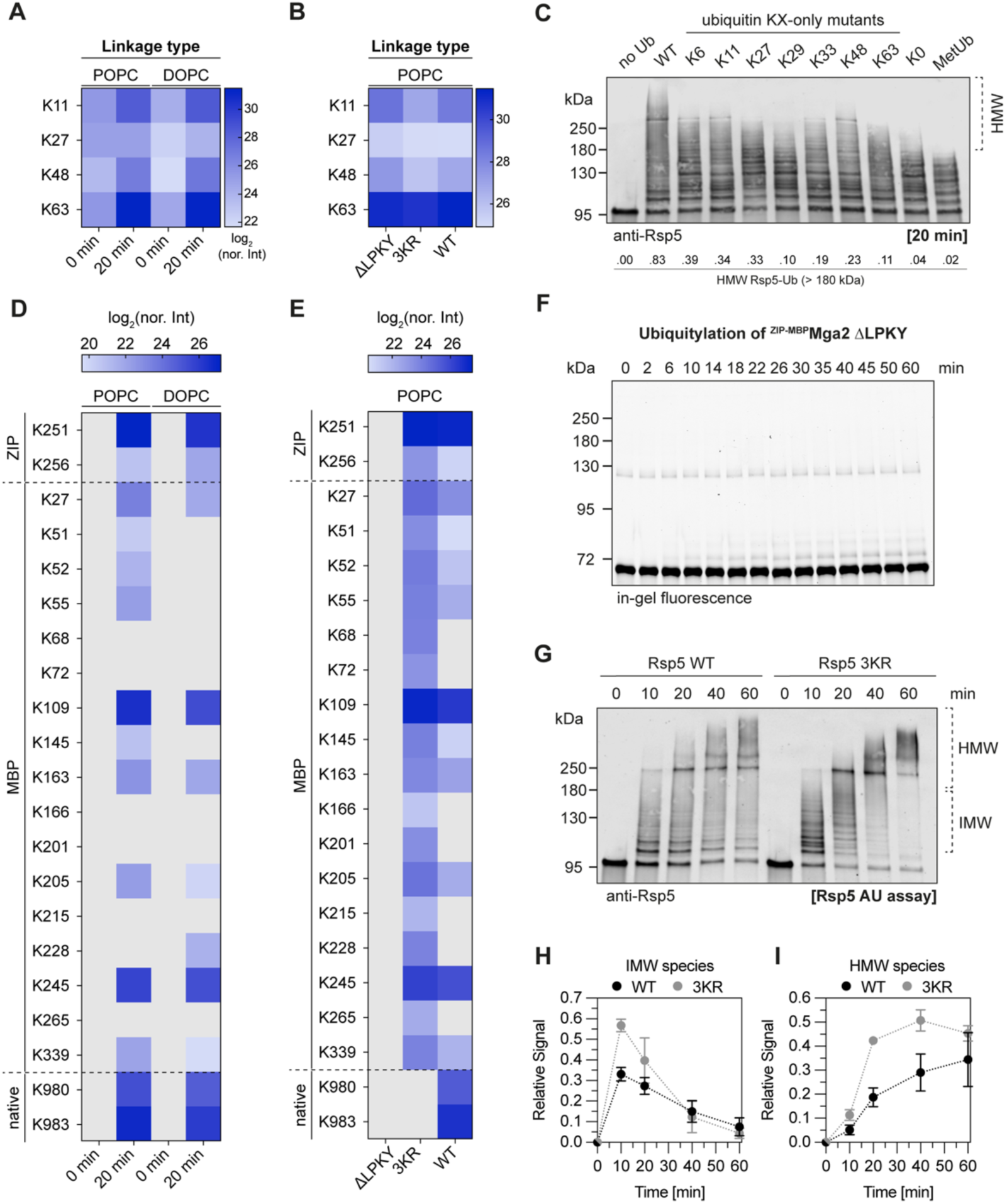
**(A)-(B)** Mass spectrometric analysis of detected ubiquitin linkage types. **(A)** Comparing ^ZIP-MBP^Mga2 ubiquitylation in POPC and DOPC membranes. **(B)** Assays performed with ^ZIP-MBP^Mga2 mutants, reconstituted into POPC membranes. ΔLPKY: deletion of the Rsp5 binding motif from Mga2. 3KR: triple lysine-to-arginine substitution of Mga2’s native lysines (K980R, K983R, K985R). **(C)** Rsp5 autoubiquitylation assays performed with ubiquitin mutants. Assay components: 70 nM E1, 500 nM E2, 200 nM Rsp5, 15 µM ubiquitin variant, ATP. Ubiquitin KX-only mutants: constructs containing only the indicated native lysine (remaining lysines substituted with arginine). K0: lysine-less constructs with lysine-to-arginine substitution of all seven lysines. MetUb: Octa-dimethyl-ubiquitin (bovine, Enzo Life Science). All seven lysines and the N-terminal amine group are methylated. Reactions were incubated at 30°C for 20 min. Samples containing 83 ng of Rsp5 were analyzed by SDS-PAGE (gels: 7.5% Mini-PROTEAN^®^ TGX™) and immunoblotting. Primary antibody: anti-Rsp5 serum, 1:1000 (rabbit, polyclonal). Secondary antibody: IRDye^®^ 800 CW anti-rabbit, 1:15000 (goat). Blots were scanned using a LI-COR Odyssey^®^ scanner at 700 and 800 nm (84 µm resolution). Signal intensities of high molecular weight Rsp5-Ub species were quantified, background-corrected using control lanes (no Ub), and normalized to unmodified Rsp5 (no Ub control). The mean of n = 3 replicates is indicated **(D) and (E)** Overview of all ^ZIP-MBP^Mga2 ubiquitylation sites detected by mass spectrometry. The log2 of normalized intensities from n = 3 replicates is shown. **(D)** Assays performed with ^ZIP-MBP^Mga2 reconstituted into POPC or DOPC membranes. **(E)** Assays performed with ^ZIP-MBP^Mga2 mutants reconstituted into POPC membranes. **(F)** *In vitro* ubiquitylation of ATTO 590-labeled ^ZIP-MBP^Mga2 ΔLPKY after reconstitution into POPC membranes. Assay components: 70 nM E1, 500 nM E2, 200 nM Rsp5, 2 µM ^ZIP-MBP^Mga2 ΔLPKY (in POPC), 15 µM ubiquitin, ATP. Reactions were performed at 30°C.Per sample, 0.55 µg of ^ZIP-MBP^Mga2 ΔLPKY were analyzed by SDS-PAGE. In-gel fluorescence was detected using a Typhoon laser scanner (635 nm laser, Cy5 filter, 25 µm resolution, PMT voltage of 500 V). **(G)** Comparing *in vitro* autoubiquitylation of WT Rsp5 and Rsp5 3KR (K411R, K432R, K438R). Assay components: 70 nM E1, 500 nM E2, 200 nM Rsp5 (WT or 3KR mutant), 15 µM ubiquitin, ATP. Reactions were performed at 30°C. Samples containing 8.3 ng of Rsp5 were analyzed as described in (C). **(H) - (I)** Quantification of Rsp5 modification. **(H)** Intermediate molecular weight species up to 180 kDa. **(I)** High molecular weight species > 180 kDa. Signals were background-corrected using intensity measurements from time point t = 0 min and normalized to the intensity of unmodified Rsp5.

The autoubiquitylation of Rsp5, on the other hand, followed an opposite trend. When the assay was performed with ^ZIP-MBP^Mga2 in loosely packed, DOPC-based membranes, Rsp5 modification was detected on five lysine residues (K45, K78, K79, K258, and K411), but only at two (K79 and K258) and with lower intensities under the POPC condition (Figure 6C). The ubiquitylation of K411, which is known attenuate E3 ligase activity (Attali *et al*, 2017), was exclusively found under the DOPC condition (Figure 6C). These findings suggest that ^ZIP-MBP^Mga2 in loosely packed membranes can promote Rsp5 autoubiquitylation.

To assess the substrate specificity of the ubiquitylation reaction, we compared wild-type ^ZIP-MBP^Mga2 reconstituted in POPC-based liposomes with a variant lacking the LPKY motif, which is crucial for Rsp5 recruitment *in vivo* (Shcherbik *et al*, 2004) (Figure 6B, EV6E,F). Deletion of this motif led to a near-complete loss of sensor ubiquitylation, including all detectable lysine modifications (Figure 6B; EV6E,F). In contrast, Rsp5 autoubiquitylation, including the inhibitory modification of K411, was markedly enhanced (Figure 6D). These data underscore the high specificity of the reaction and establish a strict requirement of the LPKY motif for Rsp5 engagement and client ubiquitylation. In the absence of a suitable substrate, ubiquitylation occurs on Rsp5 itself, presumably causing autoinhibition.

When the three prime targets of Mga2 ubiquitylation *in vivo* (K980, K983, K985) are replaced with arginine (^ZIP-MBP^Mga2 3KR), we observed a broad redistribution of the ubiquitylations to alternative lysine residues in the ^ZIP-MBP^Mga2 construct (Figure 6B, EV6E) and reduced levels of Ub-Ub linkages (Figure EV6B). The autoubiquitylation of Rsp5 remained largely unchanged in this condition (Figure 6D). Overall, these observations suggest a preferential formation of mixed polyubiquitin chains on those lysine residues, which are targeted by Rsp5 *in vivo*. Furthermore, the detailed characterization of ubiquitylation reveals a dynamic ‘rewiring’ of preferred ubiquitylation sites, which is dependent on the appropriate recruitment of Rsp5, the accessibility of the preferred ubiquitylation sites, and the lipid environment of Mga2.

Having established that the auto-inhibitory modification of Rsp5 K411 occurred selectively in loosely packed membranes, we wanted to validate if Rsp5 autoubiquitylation might contribute to the sensitivity of the OLE pathway and its ability to distinguish between rather saturated or more unsaturated membrane environments. To this end, we generated a Rsp5 variant, in which all three reported auto-inhibitory lysine residues (K411, K432, K438) (Attali *et al*, 2017) were replaced by arginine (Rsp5 3KR), and then used it to ubiquitylate wildtype ^ZIP-MBP^Mga2 in different lipid environments. While the multiubiquitylation of Mga2 by wildtype Rsp5 was sensitive to the lipid environment (Figure 6E), the ability of the system to distinguish POPC and DOPC was lost when the Rsp5 3KR variant was used.

A deeper look into the ubiquitylation kinetics revealed that that the 3KR mutation in Rsp5 had barely any impact on Mga2 ubiquitylation in the more saturated membrane environment where Rsp5 autoubiquitylation at the inhibitory K411 does not occur (Supplementary Figure S5A-H). In unsaturated membrane environments, however, the 3KR mutation in Rsp5 facilitates Mga2 ubiquitylation in particularly at late steps in the reaction (Supplementary Figure S5I-P) leading ultimately to robust differences in the level of multiubiquitylated Mga2 between different membrane environments (Figure 6E,F). Hence, the autoubiquitylation of Rsp5 is functionally relevant and contributes to signal amplification. Overall, the detailed analysis of Rsp5 and Mga2 ubiquitylation reveals a dynamic interdependence of the E3 ligase and its client, which provides the basis for signal amplification and the exquisite sensitivity of the OLE pathway.

## Discussion

Signal amplification in biology is often achieved through multilayer processes that integrate weak and noisy input signals for robust signaling outcomes. Here, we dissect a signal amplification mechanism for decisive ubiquitylation of the lipid saturation sensor Mga2. We propose that the remarkable sensitivity of Mga2 is based on lipid packing-dependent conformational changes in the transmembrane region (Covino *et al*, 2016) that are transmitted to the site of ubiquitylation (Ballweg *et al*, 2020) and amplified by the kinetics of sensor ubiquitylation. At the heart of the mechanism lies a membrane-controlled kinetic barrier for the first ubiquitin transfer and a diversion of ubiquitylation between the sensor protein and the E3 ligase itself, which may be supported by the processivity of the reaction. In tightly packed membranes, the lipid saturation sensor is rapidly ubiquitylated, while in loosely packed membranes Rsp5 autoubiquitylation dominates, thereby limiting its E3 ligase activity and further reducing sensor ubiquitylation (Figure 6G) (Attali *et al*, 2017).

### A kinetic barrier, positive feedback, and irreversible inhibition for signal amplification

Sensitive quantification of reaction intermediates over a broad dynamic range provided the basis for a detailed characterization of the reaction kinetics. Even though we used steady-state conditions allowing repeated encounters between the E3 ligase and its client, we uncovered kinetic features similar to those in single-encounter experiments, albeit on a different time scale (Pierce *et al*, 2009; Kim & Huibregtse, 2009). Firstly, the initial ubiquitin transfer to Mga2 is markedly slower than subsequent additions, thereby suggesting a kinetic barrier that must be overcome to initiate productive multiubiquitylation. Secondly, subsequent ubiquitylations occur at many-fold higher rates, indicating positive feedback, likely because ubiquitin modifications increase the local density of lysine residues and stabilize the interaction with Rsp5 (French *et al*, 2009; Kim *et al*, 2011; Zhu *et al*, 2022). Thirdly, an irreversible inhibition at every step of the reaction is crucial to reliably fit the data.

Although the lipid environment does not modulate the interaction between Mga2 and Rsp5 (Figure EV3A,B), we observe significant differences in the first round of ubiquitylation (Figure 3D) that are further amplified in subsequent rounds (Figure 3E-K). Consistent with ’ubiquitylation zones’ proposed in the context of Rsp5-dependent endocytosis (Sardana *et al*, 2019), we propose that the lipid-regulated local density of lysine residues controls the balance between Mga2 ubiquitylation and Rsp5 autoubiquitylation. This is supported by 1) lipid-dependent conformational changes around the ubiquitin attachment site revealed by FRET (Ballweg *et al*, 2020), 2) strongly reduced multiubiquitylation of Mga2 with lysine-less ubiquitin (Figure EV2B-K), and 3) increased Rsp5 autoubiquitylation when recruited by Mga2 in loosely packed lipid membranes (Figure 4E,G).

The molecular basis of the inhibitory component remains difficult to define. It is likely more complex under steady-state conditions than in single-encounter experiments, where inhibition can be attributed to the dissociation of the processive E3 ligase from its client (Pierce *et al*, 2009). Most encounters are probably unproductive, with repeated dissociation events. Under these conditions, Rsp5 can be titrated away from poorly modified clients to more heavily ubiquitylated species, contributing to incomplete consumption of reaction intermediates (Figure 2A-I). Additional factors such as E1/E2 inactivation or aggregation of the ER ligase and/or its client may also contribute. Therefore, the rate of irreversible inhibition in our model should be considered as composite term of several contributing mechanisms.

### How does the ubiquitylation kinetics contribute to decision making?

Even modest changes in ubiquitylation efficiency (forward rate over inhibition rate) can strongly influence overall sensor ubiquitylation. This is particularly critical in the first round, which acts as the ‘gatekeeper’ for multiubiquitylation (Figure 3D,M). In tightly packed membranes, the efficiency favors ubiquitylation (1.24), whereas in loosely packed membranes if favors inhibition (0.83) (Figure 3M). Subsequent rounds further amplify these differences up to the fifth ubiquitin transfer (Figure 3E-H, M).

Importantly, discrimination between saturated and unsaturated membranes also depends on Rsp5 autoubiquitylation (Figure 6E, F). Compared to WT Rsp5, the Rsp5 triple mutant (K411R, K432R, K438R) causes increased Mga2 ubiquitylation selectively in loosely packed membranes but not in saturated ones (Supplementary Figure S5A-P). Consequently, the system loses sensitivity and the ability to faithfully distinguish saturated and unsaturated membranes (Figure 6E, F).

### On the role of DUBs in the OLE pathway

Signal amplification is functional *in vitro* even in the absence of deubiquitylating enzymes. The interplay of Rsp5 and Mga2 alone is sufficient to generate in robust differences in ubiquitylation (Figure 3). We find strong *in vivo* evidence against a major role of the Ubp2 in regulating lipid saturation under normal growth conditions (Figure 4H,I; Figure EV4E-H; Supplementary Figure S4A-C), despite its known antagonism to Rsp5-dependent ubiquitylation (Kee *et al*, 2005, 2006). Notably, these observations do not exclude a contribution of DUBs to the OLE pathway. In fact, distributive, non-productive ubiquitin modifications may be removed from the lipid saturation sensor by deubiquitylating enzymes other than Ubp2 and sharpen the transition to productive modification similar to the processes for substrate selection in the ERAD pathway and substrate ordering by the APC (Rape *et al*, 2006; Zhang *et al*, 2013). The amplification mechanism described for lipid saturation sensing is reminiscent of kinetic proofreading (Hopfield, 1974), where successive modifications must occur until the system commits to proceed via an irreversible step. Each successive modification step amplifies small differences in rates or binding affinities or rate constants. Analogously, polyubiquitylation of Mga2 is a multistep process in which rate differences are amplified, resulting in sufficiently modified species that are irreversibly mobilized for downstream signaling.

### Ubiquitylation balance between Rsp5 and its client

Several observations point toward a diversion of the ubiquitylation reaction depending on the membrane environment of Mga2 and its ability to accept ubiquitin modifications. This is best illustrated by the autoinhibitory modification of K411 in Rsp5 that only occurs upon recruitment to Mga2 in loosely packed membranes but not in saturated ones (Figure 6C). More severely, when Mga2’s LPKY motif is removed and cannot recruit Rsp5 (Shcherbik *et al*, 2004) ubiquitin is transferred exclusively to the E3 ligase itself, including K411 (Figure 6B,D). In contrast to that, removal of the ’natural’ ubiquitin attachment sites from Mga2 redirects ubiquitylation within Mga2, leading to a distorted, more distributed ubiquitylation pattern (Figure 6B, EV6E). We suspect that the structural dynamics of the Mga2 in a saturated membrane environment remains largely unaffected by the 3KR substitution, thereby still directing the ubiquitylation activity away from Rsp5. This prevents Rsp5 autoubiquitylation at K411 (Figure 6D) and keeps the E3 ligase in an un-inhibited, active state, which ultimately causes broadly distributed Mga2 ubiquitylations (Figure EV6E) and reduces ubiquitin chain formation (Figure EV6B).

A fine ubiquitylation balance between an E3 ligase and its client has also been described in the regulation of membrane fluidity in mammalian cells (Ruiz *et al*, 2019; Pilon, 2021; Volkmar *et al*, 2022). Here, the regulatory circuit is provided by the E3 ligase RNF145 and the so-called adiponectin receptor 2 (ADIPOR2), which harbors an intrinsic ceramidase activity (Vasiliauskaité-Brooks *et al*, 2017; Ruiz *et al*, 2019; Pilon, 2021). In unsaturated membranes, when the ceramidase activity is not required, RNF145 ubiquitylates the ceramidase as a signal for subsequent degradation (Volkmar *et al*, 2022). In saturated membranes, RNF145 undergoes autoubiquitylation and degradation, thereby leading to the accumulation of the ceramidase ADIPOR2 and a recovery of normal membrane fluidity through direct and indirect mechanisms (Volkmar *et al*, 2022; Ruiz *et al*, 2023). Notably, the membrane-anchored E2 UBE2j2 can act as the lipid saturation sensor in this pathway as demonstrated in recent, elegant reconstitution experiments (Vrentzou *et al*, 2025).

Unlike the regulated degradation RNF145, our findings on the OLE pathway suggest an auto-regulatory feedback loop that serves to establish an inactive, but readily activatable pool of Rsp5. This makes sense considering the many functions and clients of Rsp5, which is also involved in endocytosis, vacuolar degradation, and during cytosolic heat stress (Kaliszewski & Zoładek, 2008; Fang *et al*, 2014; Ballweg & Ernst, 2017; Sardana & Emr, 2021). While Rsp5 undergoes inhibitory autoinhibition in the absence of abundant substrates, its activity can be mobilized upon a suddenly increased demands, for example upon heat stress (Fang *et al*, 2014). If this auto-regulatory cycle indeed supports the orchestration of the many Rsp5 functions, and if this mechanism has wider relevance also for other multi-functional ubiquitin ligases remains to be tested in greater details (Kaliszewski & Zoładek, 2008).

### Summary

Signal amplification by Rsp5 is facilitated by three features: 1) distinct ubiquitylation efficiencies particularly at the first, rate-limiting step, 2) accelerated ubiquitylation establishing positive feedback (Pierce *et al*, 2009; French *et al*, 2017; Markevich *et al*, 2004; French *et al*, 2009), and 3) negative feedback via inhibitory autoubiquitylation of Rsp5, which occurs favorably when Mga2 is situated in a loosely packed membranes. Future work will explore how Rsp5 autoregulation contributes to the orchestration of its many functions.

## SUPPLEMENTARY FIGURES

**Supplementary Figure S1:**
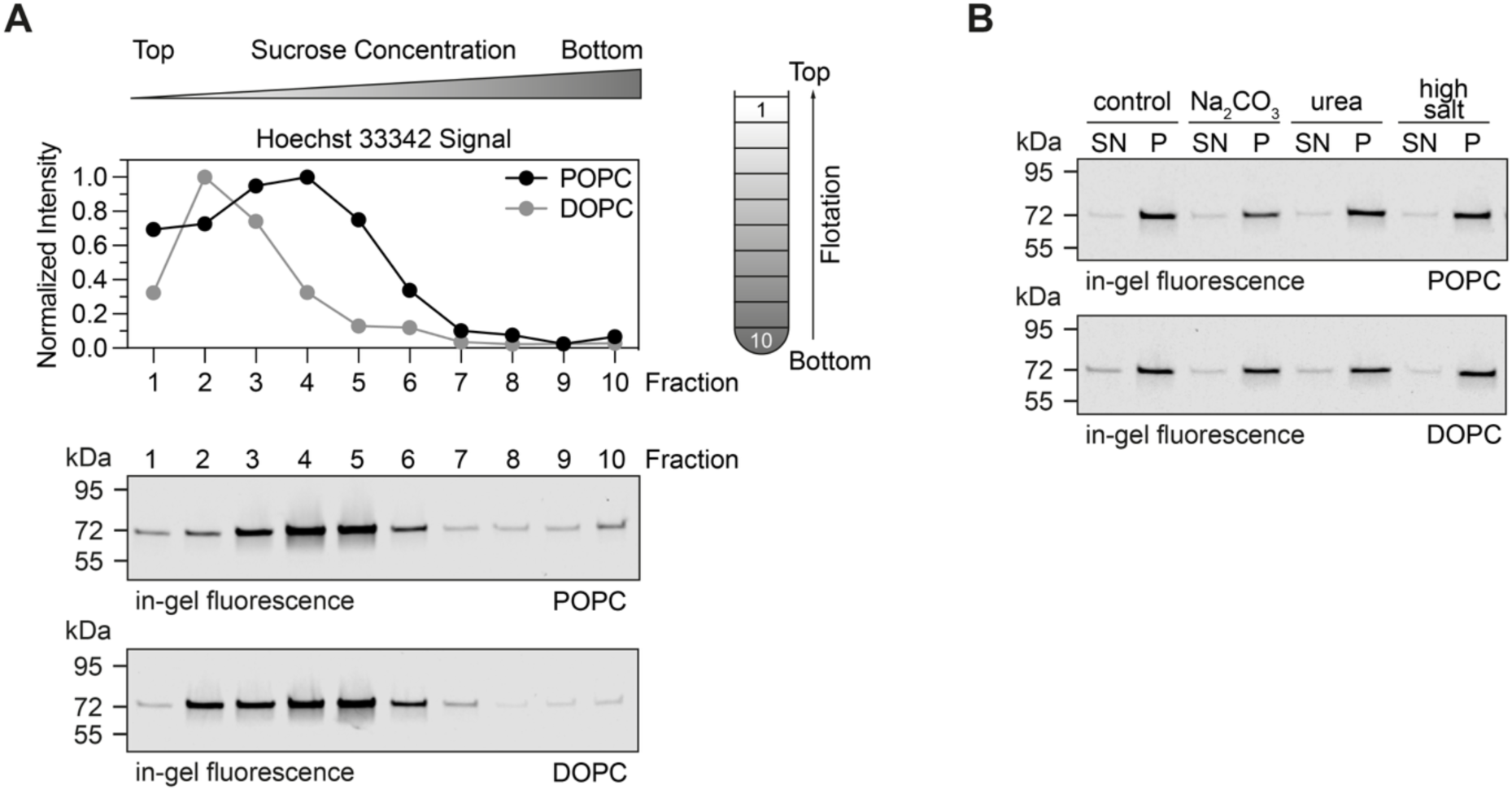
Validation of correct membrane insertion of ^ZIP-MBP^Mga2. **(A**) Incorporation of the protein into the lipid bilayer during reconstitution was tested by sucrose density gradient centrifugation. 25 µg of protein were set to 40% (w/v) sucrose and overlaid with successively decreasing sucrose concentrations (20%, 10%, 5%, 0%). The gradients were centrifuged at 100,000 x g for 18 hours and samples were collected from top (low sucrose) to bottom (high sucrose). The protein content was determined via SDS-PAGE (10 µL of sample was loaded, gels: 7.5% Mini-PROTEAN^®^ TGX™) and in-gel fluorescence scanning (Typhoon laser scanner, 488 nm laser, Cy2 filter, 25 µm resolution, 360 V PMT voltage). The lipids were tracked by staining 100 µL of each fraction with 7 µM of Hoechst 33342 (Jumpertz *et al*, 2011; Cordeiro *et al*, 2023; Halbleib *et al*, 2017). The signal was detected using a TECAN plate reader: ex. = 355 nm, em. = 459 nm, bandwidth = 20 nm. **(B)** Membrane integration and association or aggregation of ^ZIP-MBP^Mga2 were tested by alkaline carbonate extraction, urea extraction, and high salt extraction. Each condition was performed with 5 µg of protein to which buffer containing 200 mM Na_2_CO_3_, 2 M urea, or 500 mM NaCl was added. The mix was incubated for 30 min, after which proteoliposomes were separated from soluble components by ultracentrifugation (350,000 x g, 2 h, 4°C). The supernatant (SN) containing soluble components was recovered. The proteoliposome-containing pellet (P) was resuspended in a volume equal to the recovered SN (V_SN_ = V_P_). The samples of the SN and P were analyzed by SDS-PAGE (10 µL of each fraction was loaded, gels: 7.5% Mini-PROTEAN^®^ TGX™) and in-gel fluorescence scanning (Typhoon laser scanner, 488 nm laser, Cy2 filter, 25 µm resolution, 360 V PMT voltage).

**Supplementary Figure S2:**
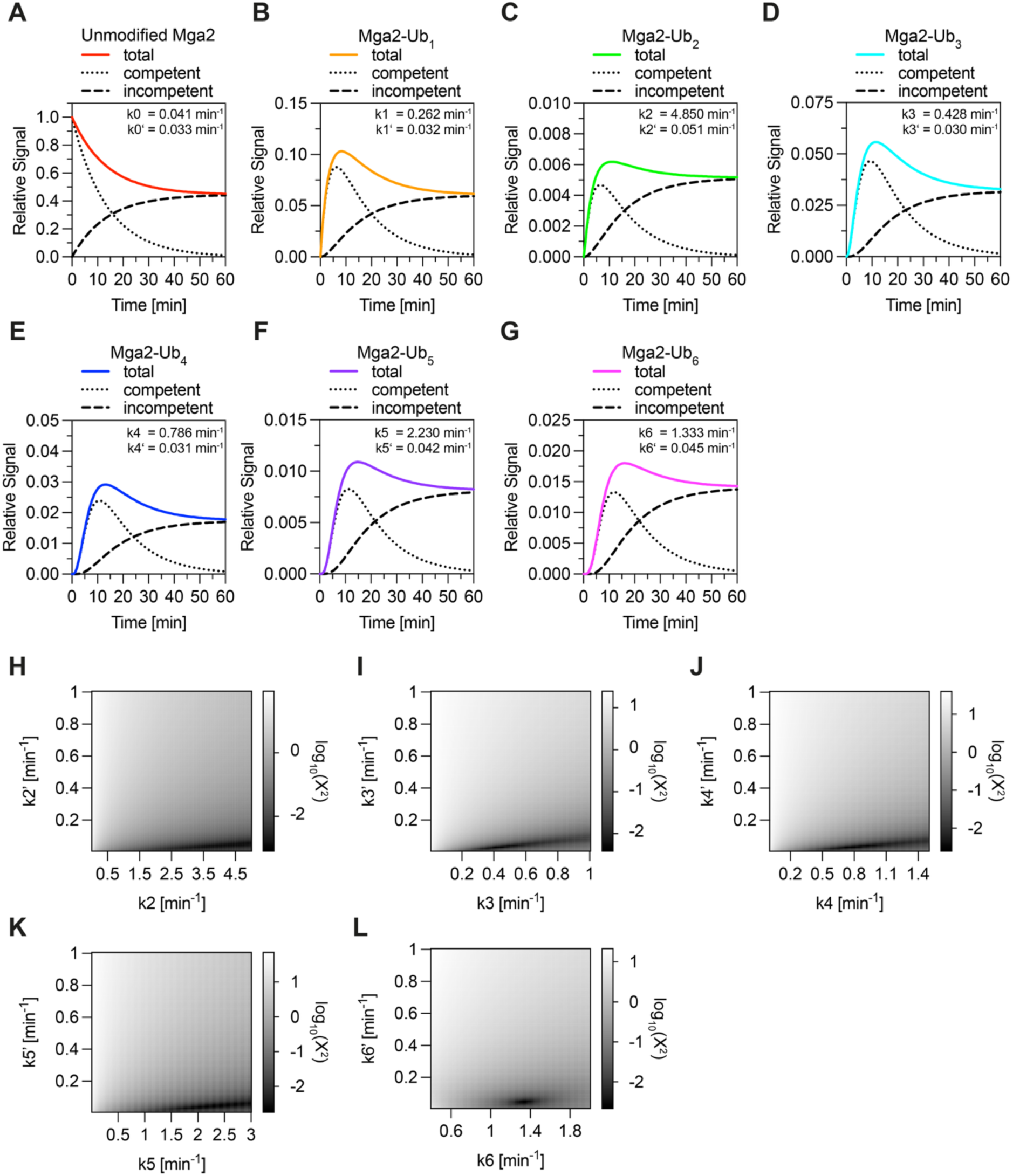
Quality assessment of fits for ^ZIP-MBP^Mga2 ubiquitylation in POPC membranes. **(A) - (G)** Contribution of ubiquitylation and inhibition to the fit. The sum of the ubiquitylated portion (dotted lines) and the inhibited pool (dashed lines) gives the overall shape of the fit (solid line). The fits are re-plotted from figure 2 and were generated by fitting the *in vitro* ubiquitylation data to the theoretical model. The optimal values for the parameters kx and kx’ are indicated on the upper right. **(H) - (L)** Heatmap analysis of the parameter space for each kx/kx’ parameter pair. The log_10_(^2^) for different kx/kx’ pairs is shown. All rates were successively increased in 0.01 min^-1^ steps. **(H)** Screened area k2/k2’: k2 from 0.01 min^-1^ to 5.0 min^-1^ and k2’ from 0.01 min^-1^ to 1.0 min^-1^. **(I)** Screened area for k3/k3’: both parameters from 0.01 min^-1^ to 1.0 min^-1^. **(J)** Screened area for k4/k4’: k4 from 0.01 min^-1^ to 1.5 min^-1^ and k4’ from 0.01 min^-1^ to 1.0 min^-1^. **(K)** Screened area for k5/k5’: k5 from 0.01 min^-1^ to 3.0 min^-1^ and k5’ from 0.01 min^-1^ to 1.0 min^-1^. **(L)** Screened area for k6/k6’: 0.4 min^-1^ to 2.0 min^-1^ and k6’ from 0.01 min^-1^ to 1.0 min^-1^.

**Supplementary Figure S3:**
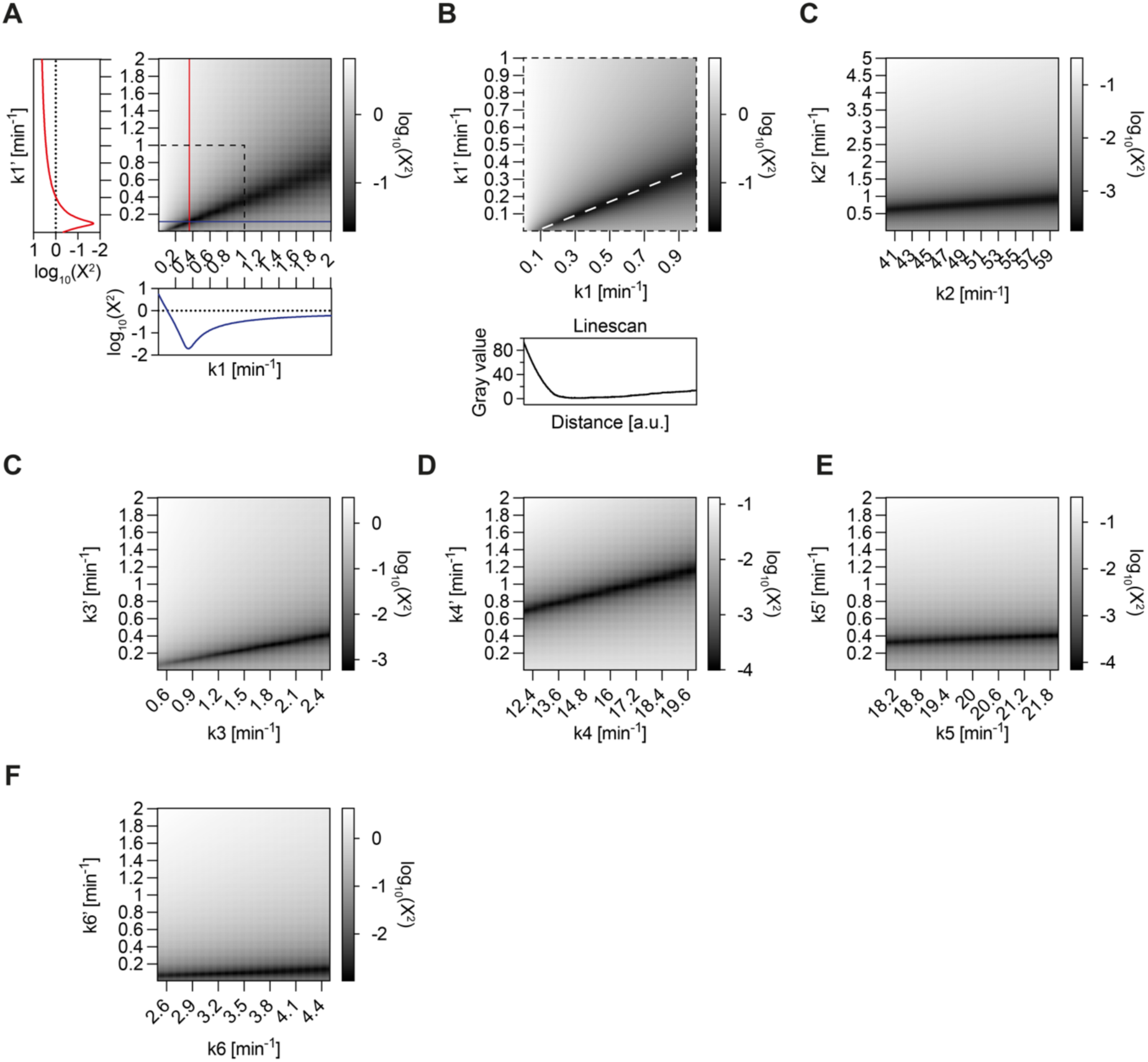
Quality assessment of fits for ^ZIP-MBP^Mga2 ubiquitylation in DOPC membranes. **(A)-(F)** Heatmap representation of log_10_(c^2^). The parameters kx and kx’ were successively increased, and the quality of the fit was evaluated. **(A)** Screened area for k1/k1’: k1 from 0.01 min^-1^ to 2.0 min^-1^ and k1’ from 0.01 min^-1^ to 1.0 min^-1^. The line scans show the effect of different complementary rates on c^2^ for a fixed rate k1 or k1’. Red curve: quality values for a fixed k1 = 0.35 min^-1^ with various k1’ values. Blue curve: quality values for a fixed k1’ = 0.11 min^-1^ with increasing values for k1. Dashed box: focus on a zoomed in area of the heatmap with increased resolution. Screened area: k1 from 0.002 min^-^1 to 1.0 min^-1^ and k1’ from 0.002 min^-2^ to 1.0 min^-1^. The rates were successively increased by 0.002 min^-1^. White dashed line: scan along the optimal values for analysis of the optimum. **(B)** Selected area for k2/k2’: k2 from 40 min^-1^ to 59.96 min^-1^ in 0.04 min^-1^ increments and k2’ from 0.01 min^-1^ to 5.0 min^-1^ in 0.01 min^-1^ steps. **(C)** Screened area for k3/k3’: k3 from 0.5 min^-1^ to 2.49 min^-1^ and k3’ from 0.01 min^-1^ to 2.0 min^-1^, both in steps of 0.01 min^-1^. **(D)** Screened area for k4/k4’: k4 from 12.0 min^-1^ to 19.96 min^-1^ in steps of 0.04 min^-1^ and k4’ from 0.01 min^-1^ to 2.0 min^-1^ with 0.01 min^-1^ steps. **(E)** Screened area for k5/k5’: k5 from 18.0 min^-1^ to 21.98 min^-1^ in steps of 0.02 min^-1^ and k5’ from 0.01 min^-1^ to 2.0 min^-1^ in 0.01 min^-1^ steps. **(F)** Screened area for k6/k6’: k6 from 2.5 min^-1^ to 4.49 min^-1^ and k6’ from 0.01 min^-1^ to 2.0 min^-1^, both in steps of 0.01 min^-1^.

**Supplementary Figure S4:**
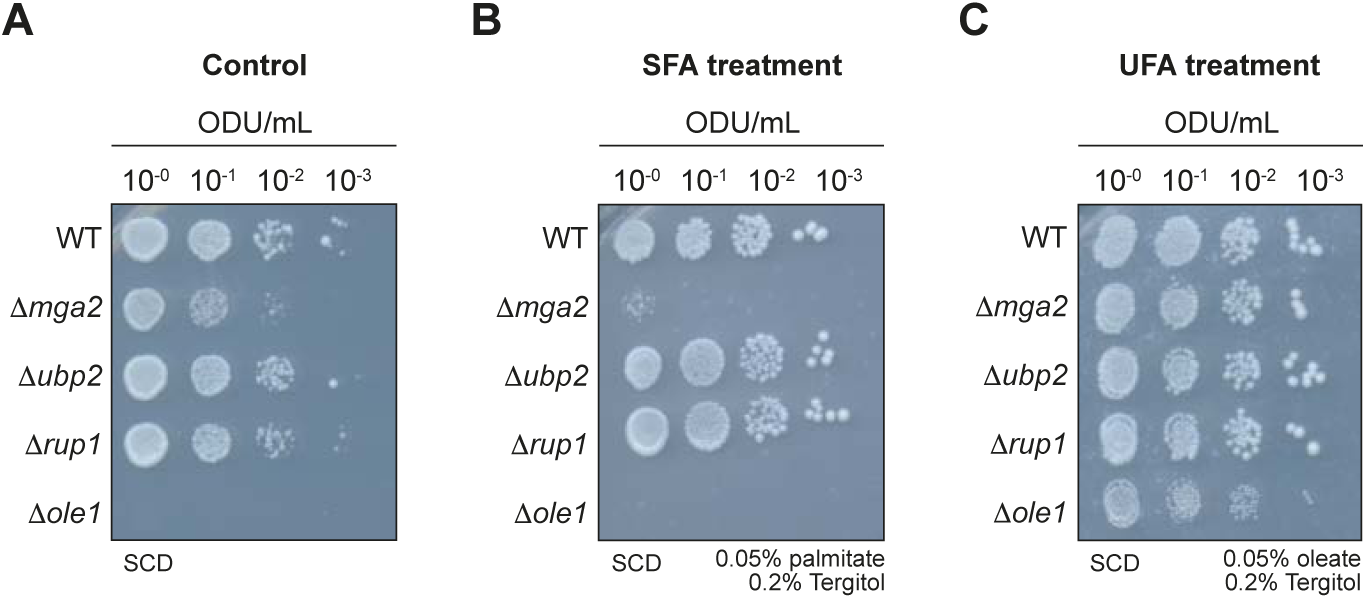
Δ*ubp2* and Δ*rup1* strains are insensitive to fatty acid supplementation. **(A) - (C)** Serial dilution growth assays on minimal media (SCD). Stationary liquid cultures of yeast were used to prepare tenfold serial dilutions, with a starting concentration of 1.0 ODU/mL. **(A)** Growth on SCD. **(B)** Growth on SCD plates containing 0.05% (w/v) sodium palmitate (w/v) and 0.2% (w/v) Tergitol^™^. **(C)** Growth on SCD plates containing 0.05% (w/v) sodium oleate (w/v) and 0.2% (w/v) Tergitol^™^.

**Supplementary Figure S5:**
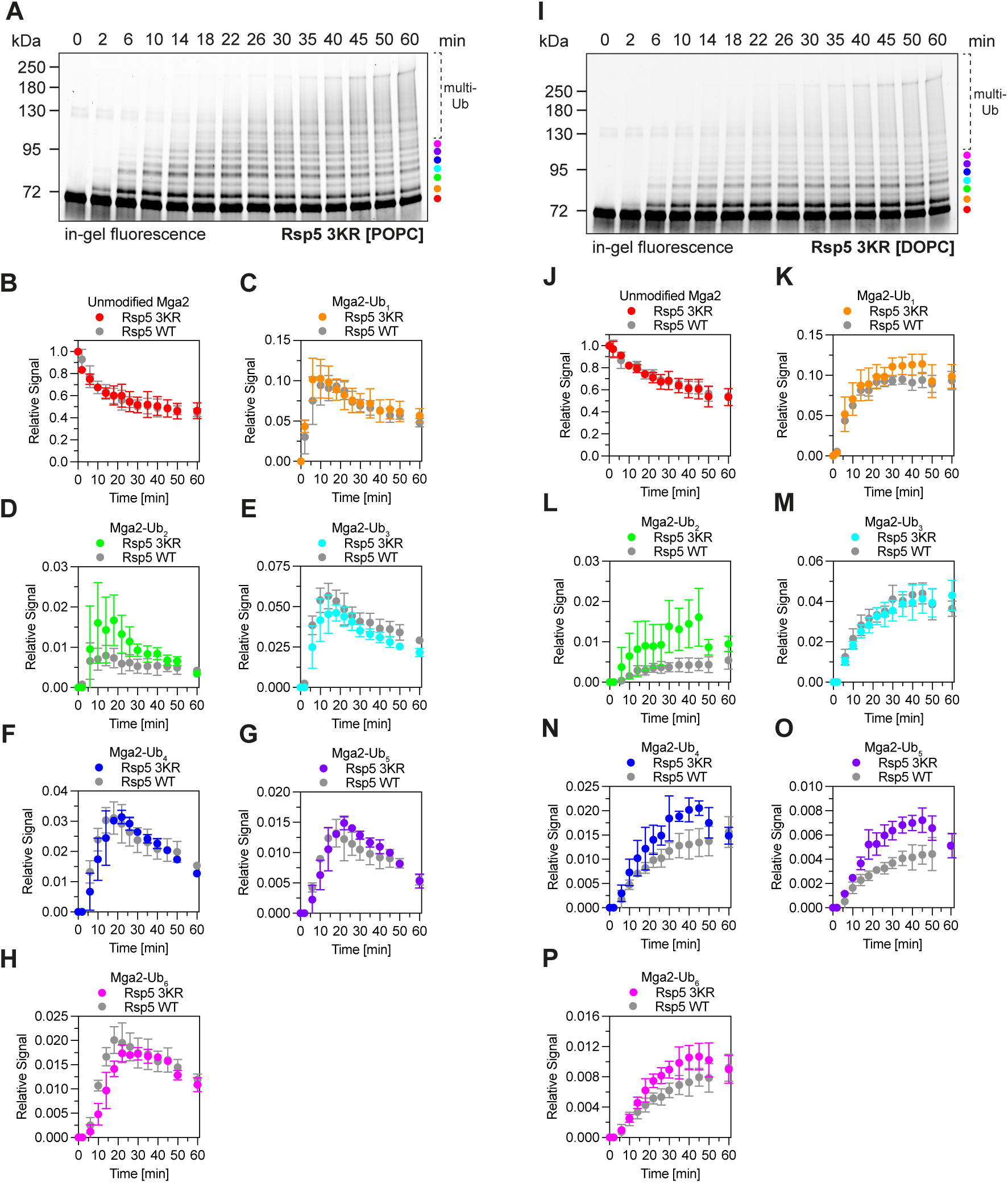
Ubiquitylation of ^ZIP-MBP^Mga2 by Rsp5 3KR. **(A)** *In vitro* ubiquitylation of ^ZIP-MBP^Mga2 in POPC membranes using Rsp5 3KR. Assay components: 70 nM E1, 500 nM E2, 200 nM Rsp5 3KR, 2 µM ZIP-MBPMga2 (in POPC), 15 µM ubiquitin, ATP. Reactions were incubated at 30°C. Samples containing 0.55 µg of ^ZIP-MBP^Mga2 were analyzed by SDS-PAGE (gels: 7.5% Mini-PROTEAN^®^ TGX™), followed by in-gel fluorescence scanning (Typhoon laser scanner, 488 nm laser, Cy2 filter, 360 V PMT, 25 µm resolution). **(B) - (H)** Quantifying the relative abundance of ^ZIP-MBP^Mga2 species. Signals were normalized to signal intensities of unmodified ^ZIP-MBP^Mga2 at time point t = 0 min. The mean and SD of n = 3 experiments are shown. **(B)** unmodified ^ZIP-MBP^Mga2, **(C)** ^ZIP-MBP^Mga2-Ub_1_, **(D)** ^ZIP-MBP^Mga2-Ub_2_, **(E)** ^ZIP-MBP^Mga2-Ub_3_, **(F)** ^ZIP-MBP^Mga2-Ub_4_, **(G)** ^ZIP-MBP^Mga2-Ub_5_, **(H)** ^ZIP-MBP^Mga2-Ub_6_. **(I)** Ubiquitylation of ^ZIP-MBP^Mga2 in DOPC membranes using Rsp5 3KR. Assays and analysis were performed as described in (A). **(J) - (P)** Quantification of ^ZIP-MBP^Mga2 ubiquitylation as described above. **(J)** unmodified ^ZIP-MBP^Mga2, **(K)** ^ZIP-MBP^Mga2-Ub_1_, **(L)** ^ZIP-MBP^Mga2-Ub_2_, **(M)** ^ZIP-MBP^Mga2-Ub_3_, **(N)** ^ZIP-MBP^Mga2-Ub_4_, **(O)** ^ZIP-MBP^Mga2-Ub_5_, **(P)** ^ZIP-MBP^Mga2-Ub_6_. The data on Mga2 ubiquitylations uses Rsp5 WT are plotted as a reference in gray and are the same data as plotted in Figure 2B-I (POPC) and Figure 3D-L (DOPC) for Rsp5 WT.

## Materials and Methods

### Molecular cloning

The oligonucleotides listed in Table 2 were used to generate plasmids with the indicated amino acid substitutions. The site-directed mutagenesis PCRs were performed using the CloneAmp^™^ HiFi PCR mix (Takara Bio). Coding sequences of Ubp2 (aa 2-1272) and Rup1 (aa 2-670) were amplified from cDNA of yeast strain BY4741 and cloned into the pGEX-4T-1 vector following a Gibson assembly strategy to create GST-tagged constructs. A TEV protease cleavage site was included between the GST-tag and the inserted genes.

**Table 1:** List of plasmids used in this study.

| Number | Vector | Construct | Source |
| --- | --- | --- | --- |
| pRE496 | pETM-m60 | 8xHis-GSG-Ubiquitin (aa 2-76) | Provided by V. Dötsch |
| pRE678 | pGEX-T2 | GST-SD-TEV-S-UBCH5B (aa 1-147) | Provided by B. Schulman (Kamadurai *et al*, 2009) |
| pRE685 | pET28 | GSS-6xHis-SSG-Thrombin-H-UBA1 (aa 1-1058) | (Carvalho *et al*, 2012) |
| pRE689 | pGEX-4T-1 | GST-SD-TEV-SGGS-Rsp5 (aa 383-809) | Provided by B. Schulman (Kamadurai *et al*, 2013) |
| pRE759 | pMALC | 2xZIP-MBP-TEV site-Mga2 (aa 951-1062) | (Ballweg *et al*, 2020) |
| pRE771 | pMALC | 2xZIP-MBP-TEV site-Mga2 (aa 951-1062, K980R, K983R, K985R) | (Ballweg *et al*, 2020) |
| pRE851 | pGEX-6P-1 | GST-SD-PreScission site-LGSPEFT-Rsp5 (aa 1-809) | Provided by Yasushi Saeki and Eri Sakata (Saeki *et al*, 2005) |
| pRE855 | pMALC | 2xZIP-MBP-TEV site-Mga2 (aa 951-1062, S1003C) | This study |
| pRE867 | pGEX-4T-1 | GST-SD-TEV site-SGGS-Rsp5 (aa 383-809, K411R, K432R, K438R) | This study |
| pRE868 | pMALC | 2xZIP-MBP-TEV site-Mga2 (aa 951-1062, $\Delta^{967}$LPKY<sup>970</sup>, S1003C) | This study |
| pRE869 | pMALC | 2xZIP-MBP-TEV site-Mga2 (aa 951-1062, K980R, K983R, K985R, S1003C) | This study |
| pRE870 | pGEX-4T-1 | GST-SD-TEV-SGGS-Ubp2 (aa 2-1272) | This study |
| pRE871 | pGEX-4T-1 | GST-SD-TEV site-SGGS-Rup1 (aa 2-670) | This study |
| pRE872 | pGEX-6P-1 | GST-SD-PreScissionsite-LGSPEFT-Rsp5 (aa 1-809_K411R_K432R_K438R) | This study |

**Table 2:**
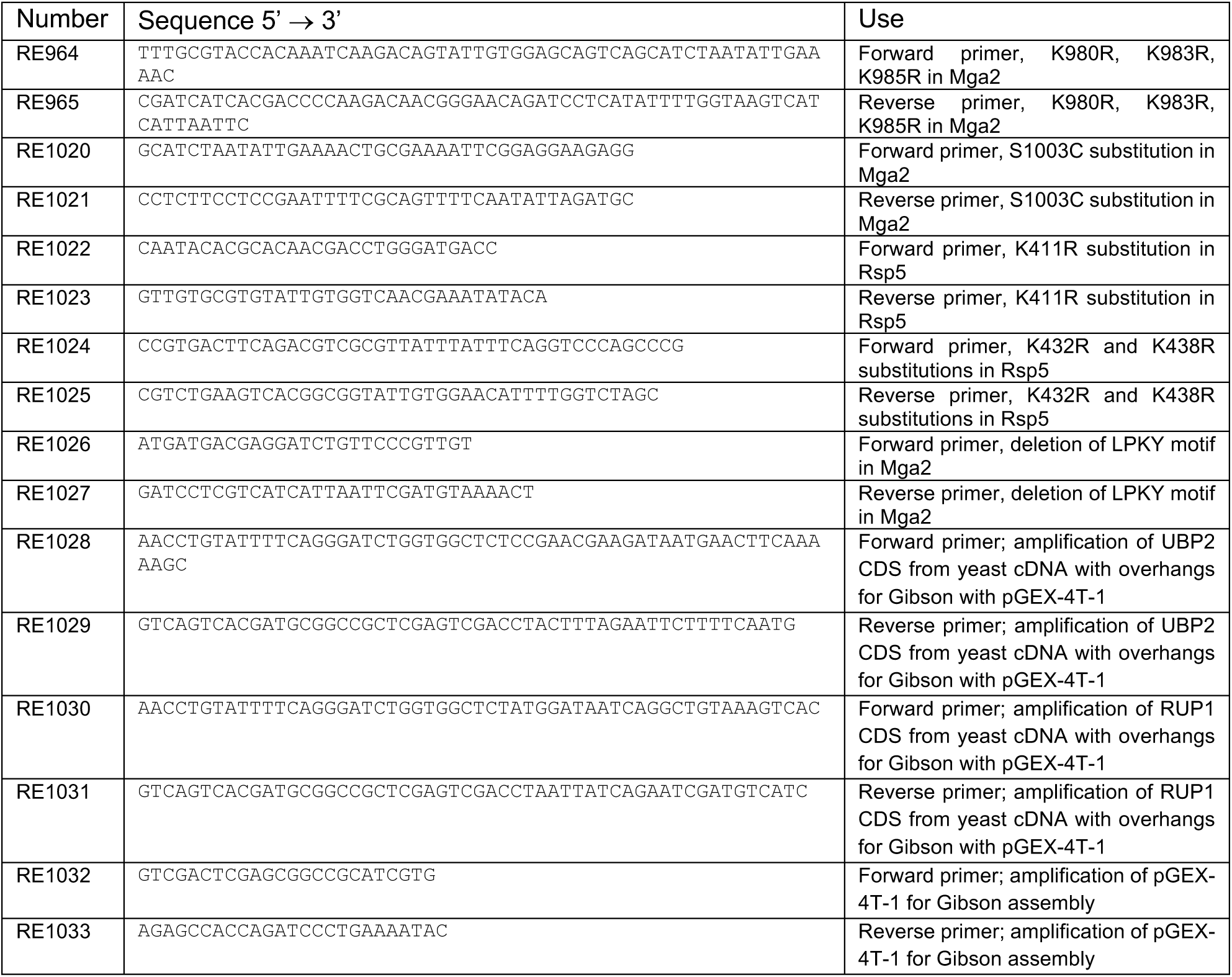
List of oligonucleotides used in this study.

### Heterologous expression of recombinant proteins

For heterologous expression of all recombinant proteins mentioned in this work, chemically competent BL21 Star (DE3) pLysS or CodonPlus-RIL (DE3) *E. coli* were transformed with the corresponding expression vectors as indicated in Table 3.

**Table 3:** List of plasmids and bacterial strains used for the heterologous production of proteins.

| Plasmid | Construct (ORF) | Bacterial strain |
| --- | --- | --- |
| pRE496 | 8xHis-GSG-Ubiquitin (aa 2-76) | BL21 Star (DE3) pLysS |
| pRE678 | GST-SD-TEV-S-UBCH5B (aa 1-147) | CodonPlus-RIL (DE3) |
| pRE685 | GSS-6xHis-SSG-Thrombin-H-UBA1 (aa 1-1058) | CodonPlus-RIL (DE3) |
| pRE689 | GST-SD-TEV-SGGS-Rsp5 (aa 383-809) | CodonPlus-RIL (DE3) |
| pRE851 | GST-SD-PreScission site-LGSPEFT-Rsp5 (aa 1-809) | CodonPlus-RIL (DE3) |
| pRE855 | 2xZIP-MBP-TEV site-Mga2 (aa 951-1062, S1003C) | BL21 Star (DE3) pLysS |
| pRE867 | GST-SD-TEV site-SGGS-Rsp5 (aa 383-809, K411R, K432R, K438R) | CodonPlus-RIL (DE3) |
| pRE868 | 2xZIP-MBP-TEV site-Mga2 (aa 951-1062, $\Delta^{967}$LPKY<sup>970</sup>, S1003C) | BL21 Star (DE3) pLysS |
| pRE869 | 2xZIP-MBP-TEV site-Mga2 (aa 951-1062, K980R, K983R, K985R, S1003C) | BL21 Star (DE3) pLysS |
| pRE870 | GST-SD-TEV-SGGS-Ubp2 (aa 2-1272) | CodonPlus-RIL (DE3) |
| pRE871 | GST-SD-TEV site-SGGS-Rup1 (aa 2-670) | CodonPlus-RIL (DE3) |
| pRE872 | GST-SD-PreScissionsite-LGSPEFT-Rsp5<br>(aa 1-809_K411R_K432R_K438R) | CodonPlus-RIL (DE3) |

The GST-tagged Rsp5 constructs, 6xHis-tagged Uba1 (E1 enzyme), GST-tagged UbcH5B (E2 enzyme), GST-tagged Ubp2, and GST-tagged Rup1 were expressed in *E. coli* cultivated in ZYM-5052-autoinduction medium (1% (w/v) tryptone, 0.5% (w/v) yeast extract, 25 mM Na_2_HPO_4_, 25 mM KH_2_PO_4_, 50 mM NH_4_Cl, 5 mM Na_2_SO_4_, 2 mM MgSO_4_, 0.5% (w/v) glycerol, 0.05% (w/v) glucose, 0.2% (w/v) α-lactose, 0.02% (v/v) 1000x trace metals) (Studier, 2005). An overnight pre-culture of bacteria cultivated in LB medium was used to inoculate a main culture (autoinduction medium) to an optical density of 0.05 OD_600_ units (ODU). The bacteria were cultivated at 37°C until an OD_600_ of 1.0 ODU was reached. The cultures were switched to incubation at 18°C for 18-20 hours to allow optimal protein synthesis. The bacteria were harvested by centrifugation (5000 x g, 30 min, 4°C) and washed with cold PBS (137 mM NaCl, 2.7 mM KCl, 10 mM Na_2_HPO_4_, 1.8 mM KH_2_PO_4_, pH 7.4). The cell pellets were stored at -20°C. For the expression of ubiquitin constructs, LB medium was used for main cultures. The ^ZIP-MBP^Mga2 constructs were cultivated in a main culture of LB medium with 0.2% (w/v) glucose. In both cases, main cultures were inoculated to an optical density of 0.05 ODU using an overnight pre-culture. The cultures were incubated at 37°C (220 rpm shaking) and gene expression was induced by addition of 0.3 mM IPTG as the cultures reached an optical density of 0.6 ODU. After three hours of expression, the bacteria were harvested by centrifugation (5000 x g, 30 min, 4°C), washed with cold PBS (pH 7.4), and stored at -20°C.

### Affinity purification of GST fusion constructs

Constructs of yeast Rsp5 (E3 enzyme), Ubp2 (DUB), Rup1, and human UbcH5B (E2 enzyme) contain an N-terminal GST-tag. Affinity purification was performed at 4°C or on ice. Frozen cell pellets were thawed and resuspended in Buffer 1 (50 mM Tris-HCl pH 8.0, 300 mM NaCl, 10% (w/v) glycerol, 0.2% (v/v) Tergitol™ Type NP-40 (Sigma-Aldrich), 1 mM DTT, 0.01% (v/v) Benzonase^®^ nuclease (Sigma-Aldrich), 0.1% (w/v) chymostatin, 0.1% (w/v) antipain, 0.1% (w/v) pepstatin A). The resuspended cells were lysed by sonication. Insoluble cell debris was removed by ultracentrifugation (100,000 x g, 30 min, 4°C). The supernatant was mixed with 3.0 mL of glutathione Sepharose™ 4B resin (Cytiva) per liter of bacterial culture. The matrix was equilibrated with water and Buffer 2 (50 mM Tris-HCl pH 8.0, 300 mM NaCl, 10% (w/v) glycerol, 1 mM DTT). The supernatant-resin suspension was incubated at 4°C for 30 min, followed by transfer to gravity columns. The resin was washed with 3x 10 column volumes (CVs) of Buffer 2. The GST-tagged protein was eluted in 3x 3.0 mL of Buffer 3 (50 mM Tris-HCl pH 8.0, 300 mM NaCl, 10% (w/v) glycerol, 1 mM DTT, 10 mM reduced glutathione) after 5 min of incubation. The eluate was collected in 1 mL fractions. The protein concentration was estimated using the absorption at 280 nm, the theoretical molar extinction coefficient, and the molecular weight of the protein. For GST-E2 preparations, the affinity-purified protein was used for size exclusion chromatography.

The GST-tag was removed from Rsp5, Ubp2, and Rup1 constructs by protease cleavage unless otherwise specified. To this end, 1 µg of TEV (short Rsp5 constructs) or PreScission^®^ protease (full-length Rsp5 constructs and Ubp2 and Rup1) was used per 100 µg of recombinant protein. Cleavage was allowed for 20 hours at 4°C with constant agitation. Then, the protease-protein mix was incubated with 500 µL of washed glutathione Sepharose™ 4B and 500 µL Ni-NTA resin for 60 min at 4°C to remove any uncleaved proteins and the proteases, respectively. The resin material was removed from the solution using gravity columns. The protein concentration was determined as described. Affinity-purified proteins were further purified by size exclusion chromatography.

### Purification and fluorescence labeling of ^ZIP-MBP^Mga2 constructs

Cell pellets were resuspended in detergent-containing Buffer 4 (25 mM HEPES pH 7.4, 150 mM NaCl, 50 mM n-octyl-D-glucopyranoside (OG), 10 mM TCEP, 1 mM EDTA, 0.01% (v/v) Benzonase^®^ nuclease (Sigma-Aldrich), 0.1% (w/v) chymostatin, 0.1% (w/v) antipain, 0.1% (w/v) pepstatin A) and lysed with ultrasound. The lysate was incubated at 4°C for 60 min with moderate agitation to allow solubilization of the membrane proteins. The insoluble components were removed by ultracentrifugation (100,000 x g, 30 min, 4°C), and the supernatant was recovered. For affinity purification, 3.0 mL of amylose resin (NEB) per liter of bacterial cultures were washed with water and Buffer 5 (25 mM HEPES pH 7.4, 150 mM NaCl, 50 mM OG, 1 mM EDTA). The recovered supernatant of the lysate was mixed with the washed resin. The supernatant-resin suspension was incubated at 4°C for 60 min with slight agitation. The suspension was transferred to gravity columns and washed with 3x 10 CVs of Buffer 5 to remove the reducing agent. The resin-bound protein was used for covalent labeling of the cysteine with a maleimide-coupled fluorophore. To this end, 3.0 mL of ATTO 488- or ATTO 590-maleimide solution (250 µM of dye in Buffer 5) were added to the gravity column and mixed with the resin-bound protein. The column was sealed and incubated at 4°C with constant, mild agitation for 18 - 20 hours. After labeling, the resin was washed with 30 CVs of Buffer 5. The labeled protein was eluted from the resin by incubating 3x 3.0 mL of Buffer 6 (25 mM HEPES pH 7.4, 150 mM NaCl, 50 mM OG, 1 mM EDTA, 10 mM maltose) for 5 min, followed by collection of 1.0 mL eluate fractions. The protein concentration and the labeling efficiency were determined for each fraction. To this end, the absorbance at 280 nm and the maximal absorption of ATTO 488 (λ_abs_ = 500 nm, λ_ems_ = 520 nm) or ATTO 590 ((λ_abs_ = 593 nm, λ_ems_ = 622 nm) were measured. The corrected protein concentration and the labeling efficiency were calculated using the protein-specific molecular weight and molar extinction coefficient and the dye-specific parameters according to the equations and specifications given by ATTO-TEC in the dye’s manual. The affinity fractions with the highest protein concentration and best labeling efficiency (>75%) were pooled and further purified by size exclusion chromatography.

### Purification of His-tagged ubiquitin constructs

Cell pellets were resuspended in Buffer 7 (50 mM HEPES pH 7.4, 250 mM NaCl, 20 mM imidazole, 0.01% (v/v) Benzonase^®^ nuclease (Sigma-Aldrich), 0.1% (w/v) chymostatin, 0.1% (w/v) antipain, 0.1% (w/v) pepstatin A) and lysed with ultrasound. The crude lysate was centrifuged (100,000 x g, 30 min, 4°C) to remove the cell debris, and the supernatant was recovered. For each liter of bacterial culture, 3.0 mL of Ni-NTA agarose (Qiagen) were equilibrated with water and Buffer 8 (50 mM HEPES pH 7.4, 250 mM NaCl, 20 mM imidazole). The equilibrated resin was mixed with the recovered supernatant. The supernatant-resin mix was incubated at 4°C for 30 min with slight agitation. The suspension was transferred to gravity columns, followed by washing with 30 CVs of Buffer 8. The resin-bound, His-tagged proteins were eluted by incubating 3x 3.0 mL of Buffer 9 (50 mM HEPES pH 7.4, 250 mM NaCl, 400 mM imidazole) for 5 min. The protein concentration was estimated using the absorbance at 280 nm and the protein-specific molar extinction coefficient (χ = 1490 M^-1^*cm^-1^) and molecular weight (MW = 9863.2 g/mol). The affinity-purified protein was further purified by size exclusion chromatography.

### Purification of His-tagged mouse E1

The protocol for affinity purification of the mouse E1 enzyme was adapted from published protocols (Carvalho *et al*, 2012). In brief, cells were resuspended in Buffer 10 (50 mM Tris-HCl pH 8.0, 150 mM NaCl, 0.15 (w/v) Triton X-100, 20 mM imidazole pH 8.0, 1 mM EDTA, 0.01% (v/v) Benzonase^®^ nuclease (Sigma-Aldrich), 0.1% (w/v) chymostatin, 0.1% (w/v) antipain, 0.1% (w/v) pepstatin A, 0.1% (v/v) PMSF) and lysed using ultrasound. The insoluble cell debris was removed by ultracentrifugation (100,000 x g, 30 min, 4°C), and the supernatant was collected. Per liter of bacterial culture, 3.0 mL of Ni-NTA agarose (Qiagen) were equilibrated with water and Buffer 11 (50 mM Na_2_HPO_4_, 150 mM NaCl, 20 mM imidazole pH 8.0, 1 mM DTT). The supernatant and resin were mixed and incubated at 4°C for 60 min with slight agitation. The resin-supernatant suspension was transferred to gravity columns, and the resin was washed with 30 CVs of Buffer 11. The bound proteins were eluted after 5 min of incubation with 3x 3.0 mL of Buffer 12 (50 mM Na_2_HPO_4_, 150 mM NaCl, 400 mM imidazole pH 8.0, 1 mM DTT). The protein concentration was determined as described. Size exclusion chromatography was performed for further purification.

### Size exclusion chromatography (SEC)

SEC was performed on ÄKTA pure 25 L chromatography systems (GE Healthcare) using a Superdex^®^ 200 Increase 10/300 GL column. For SEC with the E1, E2, and Rsp5 constructs, the column was equilibrated with detergent-free Buffer 13 (25 mM HEPES pH 7.4, 150 mM NaCl, 1 mM TCEP). For runs with the Mga2 construct, detergent-containing Buffer 14 (25 mM HEPES pH 7.4, 150 mM NaCl, 1 mM EDTA, 50 mM OG) was used. Prior to loading, the affinity-purified protein was concentrated to 600 µL using Vivaspin^®^ 20 spin concentrators (Sartorius) with the appropriate molecular weight cut-off. The concentrated protein solution was applied using a 500 µL loop. Each run was performed with a flow rate of 0.5 mL/min. Eluate fractions were collected after the void volume (8.0 mL). The protein content of each fraction was estimated using the A_280_ and the molar extinction coefficient. The labeling efficiency and protein concentration for fluorescently labeled ^ZIP-MBP^Mga2 were calculated as described. The protein was concentrated, and the glycerol content was adjusted to 20% (w/v) using glycerol stocks (80% (w/v)), prepared in the respective SEC buffers (Buffers 13 or 14). The protein concentration was set to the desired final concentration using 20% (w/v) glycerol (in Buffers 13 or 14). The protein solutions were snap-frozen in liquid nitrogen for storage at -80°C.

### Preparation of multilamellar vesicles

Multilamellar vesicles were prepared as 10 mM stocks from 18:1 (Δ9-*cis*) phosphatidylcholine (DOPC) and 16:0-18:1(Δ9-*cis*) phosphatidylcholine (POPC) (Avanti Polar Lipids). POPC and DOPC stocks (25 mg/mL) in chloroform were mixed in solvent-resistant 2.0 mL Eppendorf tubes to yield 10 µmol of lipids with a desired molar ratio of saturated and unsaturated lipid acyl chains. Chloroform was evaporated at 60°C under a constant stream of nitrogen until the formation of a lipid film. Remaining chloroform was removed using vacuum (2 - 4 mbar) for 1 hour at room temperature. Lipids were rehydrated with 1.0 mL Buffer 15 (25 mM HEPES pH 7.4, 150 mM NaCl, 5% (w/v) glycerol), followed by an incubation at 60°C with constant agitation (1200 rpm). The resulting multilamellar liposomes were sonicated for 20 min at 60°C in a water bath (VWR^®^ ultrasonic cleaner THD, power setting 9). The multilamellar vesicles were rapidly frozen with liquid nitrogen in aliquots at a lipid concentration of 10 mM.

### Reconstitution of ^ZIP-MBP^Mga2 constructs into liposomes

^ZIP-MBP^Mga2 was reconstituted into defined lipid environments with a protein:lipid ratio of 1:8000. Reconstitution mixes were prepared in the following order to limit aggregation of ^ZIP-MBP^Mga2. First, 10 mM stocks of multilamellar vesicles were solubilized with 40 mM of n-octyl-D-glucopyranoside (OG) by adding Buffer 16 (25 mM HEPES pH 7.4, 150 mM NaCl, 5% (w/v) glycerol, 20% (w/v) OG). The mix was incubated at 4°C for 10 min with mild rotation. Second, Buffer 16 and Buffer 17 (25 mM HEPES pH 7.4, 150 mM NaCl, 50 mM OG, 1 mM EDTA, 20% (w/v) glycerol) were added to reach an OG concentration of 23.5 mM. The mix was incubated at 4°C with slight agitation. Third, fluorescently labeled ^ZIP-MBP^Mga2 (0.1 mg/mL) was added last to reach a final OG concentration of 25.5 mM and a protein:lipid ratio of 1:8000. The mix was incubated at 4°C for 10 min with constant rotation. For detergent removal, 9.0 mL of reconstitution mix were dialyzed against 1.0 L of Buffer 18 (25 mM HEPES pH 7.4, 150 mM NaCl, 5% (w/v) glycerol, 1 mM EDTA). A total of 3.0 mL of the prepared reconstitution mix were transferred to Slide-A-Lyzer™ G2 dialysis cassettes with a molecular weight cut-off of 10 kDa (Thermo Scientific). The dialysis was performed at 4°C with constant mixing of the dialysis buffer in a total of four steps. First, the reconstitution mixes were dialyzed for 1 hour against 1 L of Buffer 18 containing 400 mg of Bio-Beads^®^ SM-2 resin (Bio-Rad) to provide a sink for detergent molecules. Then, two steps of dialysis against 1 L of fresh Buffer 18 for 1 hour followed. Fourth, the cassettes were placed in 1 L of Buffer 18 containing 800 mg of Bio-Beads^®^ SM-2 resin and incubated for 16-18 hours. The reconstitution reactions containing the proteoliposomes were recovered from the cassettes. The proteoliposomes were diluted 1:5 with Buffer 19 (20 mM HEPES pH 7.4, 75 mM NaCl) and harvested by centrifugation (257,000 x g at 4°C, 20 h). The pelleted proteoliposomes were resuspended in Buffer 15 to reach a final concentration of ^ZIP-MBP^Mga2 of 4.0 µM. The proteoliposomes were snap-frozen in liquid nitrogen and stored at -80°C.

### Determination of protein recovery after dialysis

Samples of the reconstitution mixes were taken before and after the dialysis procedure (see section above). OG was added to 50 µL of these samples to reach a final OG concentration of 50 mM. The final volume was set to 150 µL by adding Buffer 15. The fluorescence intensity of the samples was determined by scanning in a TECAN plate reader (ex. = 485 nm, em. = 535 nm, bandwidth = 20 nm). To this end, 100 µL of each sample were transferred to a 96-well plate (black, flat bottom, chimney well, non-binding, Greiner Bio-One). The fluorescence intensity of each sample was corrected to a background sample (50 mM OG in Buffer 16). Dilution or concentration of the fluorescence signal was accounted for by considering the total volume of the reconstitution mix before and after dialysis. Protein recovery after dialysis was normalized to the fluorescence signal of the reconstitution mix prior to dialysis.

### Preparation of large unilamellar vesicles by extrusion

Large unilamellar vesicles (LUVs) were generated from MLVs (1 mM final lipid concentration) by extrusion as previously described (MacDonald *et al*, 1991). In brief, MLVs were passed through the extruder with a 100 nm filter 21 times to create LUVs. The LUVs were diluted 1:6 with Buffer 19 and harvested by centrifugation (500,000 x g, 16 h, 4°C). The pelleted LUVs were resuspended in a small volume to increase the lipid concentration to 10 mM. The LUVs were briefly stored at 4°C and quickly used after preparation.

### Urea, carbonate, and high salt extraction of membrane associated proteins

20 µL of the proteoliposomes (4 µM protein concentration) were mixed with an equal volume of carbonate buffer (Buffer 20: 20 mM HEPES pH 7.4, 75 mM NaCl, 200 mM Na_2_CO_3_ pH 11), urea buffer (Buffer 21: 20 mM HEPES pH 7.4, 75 mM NaCl, 2 M urea), high salt buffer (Buffer 22: 20 mM HEPES pH 7.4, 500 mM NaCl) or Buffer 19 (20 mM HEPES pH 7.4, 75 mM NaCl). The mixes were then incubated at room temperature for 30 min. The mixes were diluted 25-fold with Buffer 19 followed by a centrifugation step with 350,000 x g at 4°C for 2 hours. The supernatant was recovered. The pellet was resuspended in a volume equal to the supernatant. Samples of each fraction were then mixed with 5x membrane sample buffer, boiled at 95°C for 5 min and analyzed via SDS-PAGE (10 µL of samples were loaded) followed by in-gel fluorescence scanning (Typhoon Laser Scanner, Cy2 Laser, 25 µm resolution, 360 V PMT voltage).

### Proteinase K protection assay

The reaction mix contained 12.5 µg of reconstituted ^ZIP-MBP^Mga2 (final protein concentration 0.25 µg/µL), which was diluted from the concentrated proteoliposome stock with Buffer 15 (25 mM HEPES pH 7.4, 150 mM NaCl, 5% (w/v) glycerol). To reach complete degradation of the protein, 1% (w/v) SDS was added to one condition. Then, 1 µL of proteinase K (New England Biolabs) was added to the SDS-free and SDS-containing reactions. An untreated reaction (proteoliposomes without proteinase K or SDS) served as a control. The reactions were incubated at room temperature for 60 min. The proteinase K was inactivated by boiling at 90°C for 10 min and addition of 0.1 mM PMSF. The inactivation procedure was also performed for the control condition. Then, 5x membrane sample buffer was added to the reactions, followed by an incubation at 95°C for 5 min. The samples were analyzed by SDS-PAGE and in-gel fluorescence detection to trace the degradation of the fluorescently labeled ^ZIP-MBP^Mga2 construct (Typhoon Laser Scanner, Cy2 Laser, 25 µm resolution, 360 V PMT voltage).

### Dynamic light scattering measurements

Dynamic light scattering (DLS) was performed on a Zetasizer Nano-S (Malvern Panalytical). For measurements, 50 µL of proteoliposomes (12.5 µg of ^ZIP-MBP^Mga2) were used in a ZEN2112 quartz cuvette. The temperature was allowed to equilibrate to 30°C for 2 min. The material properties were set to phospholipid-based proteoliposomes (refractive index: 1.450, absorbance: 0.001) and the buffer composition was adjusted accordingly (viscosity: 0.9298 cP, refractive index: 1.338).

### Sucrose density gradient centrifugation

The proteoliposomes with a protein content of 25 µg were adjusted to 40% (w/v) sucrose (prepared in Buffer 15) in a volume of 600 µL. The protein solution was transferred to 13.2 mL open-top thin-wall ultracentrifugation tubes. This suspension was overlaid with 2.0 mL layers of successively decreasing sucrose concentrations: 20%-10%-5%-0% (w/v) sucrose. The gradients were centrifuged in a SW41-Ti swing-out rotor at 100,000 x g for 18 hours (temperature: 4°C). The centrifuge was set to accelerate slowly and decelerate without brakes. Then, 1 mL fractions were collected from top to bottom. The protein content of each fraction was assessed by SDS-PAGE and in-gel fluorescence scanning (Typhoon Laser Scanner, Cy2 Laser, 25 µm resolution, 360 V PMT voltage).

### *In vitro* ubiquitylation assays under constant turnover conditions

*In vitro* ubiquitylation assays consisted of 70 nM E1 (6xHis-Uba1, mouse), 500 nM E2 (GST-UbcH5B, human), 200 nM E3 (Rsp5, yeast), 2 µM reconstituted fluorescently labeled ^ZIP-MBP^Mga2, and 15 µM 8xHis-ubiquitin (human) in Buffer 15 (25 mM HEPES pH 7.4, 150 mM NaCl, 5% (w/v) glycerol). Autoubiquitylation assays of Rsp5 were performed under the same conditions, but without reconstituted ^ZIP-MBP^Mga2. All components except for the 10x ATP regenerating system (10 mM ATP, 500 mM creatine phosphate, 2 mg/mL creatine phosphokinase in 25 mM HEPES pH 7.4, 150 mM NaCl, 5 mM MgCl_2_) were mixed. The ubiquitylation was started by the addition of the 10x ATP regenerating system. A master mix of the reaction was incubated at 30°C under constant shaking (300 rpm). Samples were drawn at indicated time points by mixing 15 µL of the reactions with 5 µL of 4x MSB (100 mM Tris-HCl pH 6.8, 8 M urea, 3.2% (w/v) SDS, 0.15% (v/v) bromophenol blue, 4% (v/v) glycerol, 5 mM EDTA) and incubation at 95°C for 5 min.

### Co-immunoprecipitation of ^ZIP-MBP^Mga2 and Rsp5

Magnetic Protein G-coated Dynabeads^™^ (Invitrogen) were used for immunoprecipitation of ^ZIP-MBP^Mga2. 50 µL of beads were washed three times with 400 µL of Buffer 16 (25 mM HEPES (pH 7.4), 150 mM NaCl, 5% (w/v) glycerol). Washed beads were resuspended in 100 µL of Buffer 16 containing 6.0 µL of anti-MBP antibody (murine, monoclonal, New England Biolabs). Antibodies were allowed to bind at RT for 20 min, followed by three wash steps with 400 µL Buffer 16 to remove unbound antibody. IP reactions contained 1 µM of ^ZIP-MBP^Mga2 (in DOPC or POPC membranes) and 100 nM Rsp5 in a final volume of 50 µL in Buffer 16 Detergent-solubilized controls were performed with Buffers containing 50 mM OG, and ^ZIP-MBP^Mga2 from purified protein stocks. The prepared reactions were added to the anti-MBP-loaded beads and the mix was incubated at RT for 30 min. After binding, the supernatant was collected and the beads were washed once with 100 µL Buffer 16 (Buffer 16 + 50 mM OG for detergent-solubilized samples). The bound material was eluted by adding 37.5 µL 1x MSB, 1 mM TCEP, and 1% (w/v) SDS, followed by incubation at 95°C for 5 min. All collected samples were analyzed by SDS-PAGE and in-gel fluorescence scanning and anti-Rsp5 immunoblotting.

### Antibodies

The anti-Rsp5 antibody (rabbit, polyclonal) was kindly provided by Jeffrey Brodsky. The polyclonal anti-Rsp5 serum was generated by the antibody production facility of the Department of Medical Biochemistry and Molecular Biology of Saarland University headed by Dr. Martin Jung. Rabbits were immunized with the WW-HECT domain construct of Rsp5 (aa 383-809). The anti-MBP antibody was purchased from New England Biolabs. All secondary IRDye^®^ 800CW antibodies were purchased from LI-COR Biosciences.

### SDS-PAGE and sample analysis

The samples collected in the *in vitro* ubiquitylation assays were analyzed by SDS-PAGE using 7.5% Mini-PROTEAN^®^ TGX™ or 7.5% Criterion™ TGX™ precast gels (Bio-Rad). Per well, 6 µL of sample containing 0.55 µg of ^ZIP-MBP^Mga2 was loaded. The gels were run at 180 V (Mini-PROTEAN^®^ TGX™ gels) or 200 V (Criterion™ TGX™ gels). Fluorescently labeled ^ZIP-MBP^Mga2 species were subsequently detected by in-gel fluorescence detection (ATTO 488-labeled: Cy2, laser, 25 µm resolution, photomultiplier voltage of 360 V; ATTO 590-labeled: Cy5 laser, 25 µm resolution, photomultiplier voltage of 500 V). Unlabeled Rsp5 species and the MBP were detected via immunoblotting. The proteins were blotted onto nitrocellulose membranes using the Trans-Blot^®^ Turbo™ transfer system (Bio-Rad). The membranes were blocked in Buffer 23 (5% (w/v) BSA in TBS-T (20 mM Tris pH 7.5, 137 mM NaCl, 0.2% (v/v) Tween-20)) for 30 min. The primary anti-Rsp5 serum was diluted 1:2000 in Buffer 23 and incubated with the membranes for 1 hour at room temperature or at 4°C for overnight incubation. The membranes were washed with TBS-T (20 mM Tris-HCl pH 7.5, 137 mM NaCl, 0.2% (v/v) Tween-20). The IRDye^®^ 800CW goat anti-rabbit IgG secondary antibody (LI-COR Biosciences) was prepared in Buffer 23, applied in a 1:15000 dilution, and incubated with the membranes for 1 hour at room temperature. The membranes were washed with TBS-T prior to imaging on an Odyssey^®^ CLx imaging system (LI-COR Biosciences) at both 700 and 800 nm.

### Preparation and processing of samples for mass spectrometry

*In vitro* ubiquitylation assays were performed for 20 min as described and stopped with 1 mM of N-ethyl-maleimide at indicated time points. The reaction was mixed with SDC buffer (2% sodium deoxycholate (SDC), 1 mM TCEP, 4 mM CAA, 50 mM Tris pH 8.5) and heated for 10 min at 95°C. 100 ng of trypsin and LysC were added in 50 mM Tris pH 8.5 and incubated overnight at 37°C. The digestion was stopped upon addition of 150 µL of 1% TFA in isopropanol. Peptide clean-up was performed using SDB-RPS stage tips (Sigma-Aldrich) using a wash step with 1% TFA in isopropanol and then 0.2% TFA in water. Purified peptides were eluted in 80% acetonitrile plus 1.25% ammonia and dried in a vacuum concentrator.

### Mass spectrometry data acquisition

Samples were analyzed on a Q Exactive HF coupled to an EASY-nLC 1200 (ThermoFisher Scientific) using a 35 cm long, 75 µm ID fused-silica column packed in-house with 1.9 µm C18 particles (ReproSil-Pur, Dr. Maisch) and kept at 50°C using an integrated column oven (Sonation). HPLC solvents consisted of 0.1% formic acid in water (Buffer A) and 0.1% formic acid, 80% acetonitrile in water (Buffer B). Peptides were eluted by a linear gradient from 5% to 30% B over 30 minutes, followed by a stepwise increase to 95% B in 6 minutes which was held for another 9 minutes. Full scan MS spectra (350-1650 m/z) were acquired in Profile mode at a resolution of 60,000 at m/z 200, a maximum injection time of 20 ms and an AGC target value of 3 x 10^6^. Up to 15 most intense peptides per full scan were isolated using a 1.4 m/z window and fragmented using higher energy collisional dissociation (normalized collision energy of 27). MS/MS spectra were acquired in profile mode with a resolution of 30,000, a maximum injection time of 54 ms and an AGC target value of 1 x 10^5^. Singley charged ions, ions with a charge state above 5 and ions with unassigned charge states were not considered for fragmentation.

### Mass spectrometry data analysis

MS raw data were analyzed using MaxQuant (v1.6.7.0) (Tyanova *et al*, 2016). Acquired spectra were searched against a custom database containing Mga2, Rsp5 (both NCBI txid: 1247190), Uba1 (NCBI txid: 10090), UbcH5B and ubiquitin sequences (both NCBI txid: 9606) and a collection of common contaminants using the Andromeda search engine integrated in MaxQuant (Cox *et al*, 2011). Identifications were filtered to obtain false discovery rates (FDR) below 1% for both peptide spectrum matches (PSM; minimum length of 7 amino acids). Spectra were searched with a mass tolerance of 6 ppm in MS mode, 20 ppm in HCD MS2 mode, strict trypsin specificity, and allowing up to 2 miscleavages. Carbamidomethylated cysteine was set as a fixed modification and oxidation of methionine, N-terminal protein acetylation and GlyGly on Lysine as variable modifications allowing up to 5 modifications per peptide.

### Yeast strains

| Strain | Genotype | Backgr. | Source |
| --- | --- | --- | --- |
| yRE01 | *MATa; his3Δ1; leu2Δ0; met15Δ0; ura3Δ0* | BY4741 | EUROSCARF |
| yRE06 | *MATa; his3Δ1; leu2Δ0; met15Δ0; ura3Δ0; mga2Δ::kanMX4* | BY4741 | EUROSCARF |
| yRE491 | *MATa; his3Δ1; leu2Δ0; met15Δ0; ura3Δ0; ole1Δ::kanMX4* | BY4741 | Jonas Michaelis |
| yRE757 | *MATa; his3Δ1; leu2Δ0; met15Δ0; ura3Δ0; ubp2Δ::kanMX4* | BY4741 | EUROSCARF |
| yRE801 | *MATa; his3Δ1; leu2Δ0; met15Δ0; ura3Δ0; rup1Δ::kanMX4* | BY4741 | EUROSCARF |

### Yeast spotting assays

Precultures were inoculated to an OD_600_ of 0.4 ODU/mL and incubated at 30°C for 19 h. 5 ODU were harvested and washed two times with 1 mL of ddH_2_O. The yeast were resuspended in water, setting the final concentration to 1 ODU/mL. Tenfold serial dilutions were prepared and transferred to SCD agar (1.5% (w/v)) plates using a microplate replicator. For growth on fatty acid-containing media, 0.05% (w/v) of palmitate or oleate (sodium salts) and 0.2% (w/v) Tergitol™ were added to the media. The plates were imaged after three days of incubation at 30°C.

### Sample preparation for lipidomic analyses

Single colonies were used to inoculate SCD medium to an OD_600_ of 0.4 ODU/mL, and the precultures were incubated at 30°C for 19 h. Precultures were used to inoculate 25 mL of SCD to an OD_600_ of 0.1 ODU/mL. The main cultures were incubated at 30°C (220 rpm) until an optical density of 0.8 ODU/mL was reached. 16 ODU of yeast were harvested by centrifugation, immediately placed on ice, and washed thrice with cold 155 mM ammonium bicarbonate supplemented with 10 mM sodium azide. The washed pellets were snap-frozen in liquid nitrogen. The yeast were thawed on ice and resuspended in 1 mL of 155 mM ammonium bicarbonate + 10 mM sodium azide. The resuspension was transferred to tubes containing 200 µL of 0.5 mm zirconia beads. Cells were lysed at 4°C using a DisruptorGenie bead beater for 10 min. 350 µL of the lysate were recovered and snap-frozen in liquid nitrogen for lipidomic analysis. All lipidomic analyses were performed by Lipotype GmbH (Dresden, Germany).

### Data Fitting

The model used for fitting was adapted from Pierce *et al*. (Pierce *et al*, 2009) (analytical solutions to differential equations, see below). Kinetic rate constants were estimated by χ^2^ minimization of mean normalized data. Parameter pairs were optimized through an iterative refinement procedure analogous to simulated annealing: initial search intervals covered four orders of magnitude (starting at 0.001 min⁻¹), with 1000 samples per iteration. The best-performing values defined the subsequent interval boundaries. This process was repeated 1000 times per parameter pair.

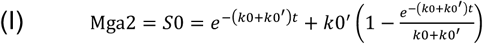

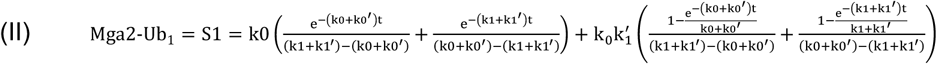

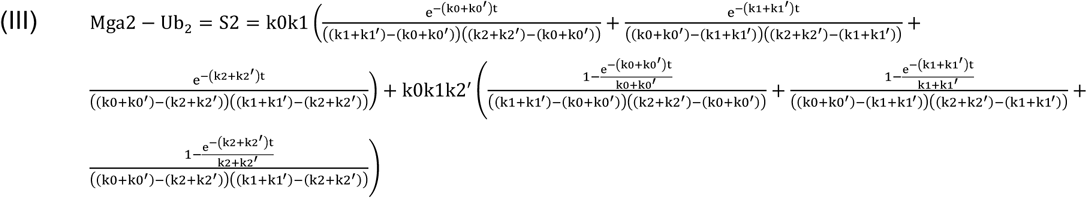

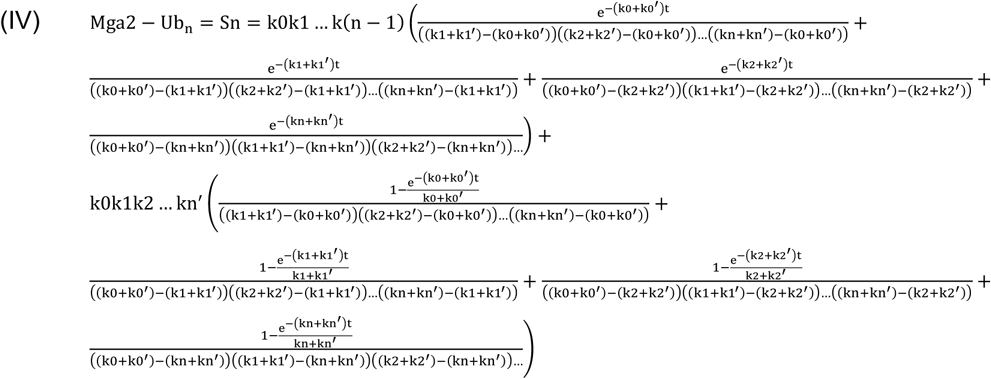

For unmodified Mga2, rates were derived assuming that abundance after 60 min is solely determined by the inhibitory rate k0’, allowing exact determination of k0 and k0’. Fit quality was quantified as weighted χ^2^, defined as the sum of squared deviations between calculated curves (y_fit_) and mean data (y_mean_), normalized by the standard deviation (SD) at each time point:

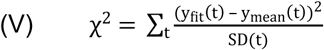

Global parameter landscapes were assessed by heat map analysis, in which parameters were varied pairwise in fixed increments and evaluated by χ^2^. All computations were implemented in C++.

## Acknowledgements

We would like to express our gratitude to Tom Rapoport, Weng Nang, Thomas Sommer, David Teis, and Alexander Stein for their critical comments and helpful suggestions. We acknowledge Heike Stumpf for excellent technical support. This work was funded by the Deutsche Forschungsgemeinschaft in the framework of the SFB1027 to HR and RE, and with an LC-MS system (easy nLC 1200, QExactive HF) used in this study (Project-ID: 259130777, SFB1177 – Selective Autophagy). Furthermore, the project was funded by the European Research Council under the European Union’s Horizon 2020 research and innovation program (grant agreement no. 866011) to RE.

